# Human *De Novo* Microproteins are Intrinsically Disordered and Contain Short Linear Motifs

**DOI:** 10.64898/2026.11.22.768061

**Authors:** Isabella Menart, Margaux Aubel, Lars Eicholt, Priyanka Madhu, Leandro Simonetti, Maria Selmer, Klara Hlouchová, Erich Bornberg-Bauer, Per Jemth, Ylva Ivarsson

## Abstract

*De novo* genes, arising from previously non-coding regions, have defied our view of gene birth and proteome evolution. Advances in ribosome profiling and comparative genomics have uncovered widespread translation of *de novo* small open reading frames (sORFs), many of which encode previously unrecognized microproteins. Despite increasing evidence demonstrating translation of *de novo* sORFs, it remains poorly understood how their encoded microproteins acquire molecular functions. Here, we selected 30 *de novo*-emerged microproteins for structural characterization and investigated their potential for interacting with canonical protein domains. Using circular dichroism and small angle X-ray scattering we found that the studied young microproteins are disordered but may adopt helical conformations in specific contexts. Additionally, we identified predicted short linear motifs (SLiMs) recognized by canonical peptide-binding domains in 29 microprotein sequences, suggesting a potential for interaction with the conserved proteome. *In vitro* experimental validation revealed interactions with relevant affinities, with some on par with canonical interactions. We also show how a microprotein SLiM can easily evolve towards higher affinities for the binding partner. Together, our findings suggest that the intrinsically disordered *de novo* microproteins can exploit SLiM-mediated molecular recognition to engage the pre-existing proteome, providing a potential mechanism for their functional integration into cellular interaction networks.

**Significance:** *De novo* gene birth from ancestrally non-coding DNA was long thought to be impossible, but increasing evidence points to the ubiquity of *de novo* protein-coding genes in all eukaryotes. While mechanisms driving *de novo* gene emergence are generally well understood, there is a substantial lack of experimental data on the products of *de novo* genes, namely *de novo* microproteins, which precludes our understanding of the emerging microproteome. To address this gap, we investigated the structures of *de novo* microproteins and their interactions with conserved protein domains. Our experimental findings support that *de novo* microproteins are predominantly disordered, and that they encode short linear binding motifs that provide an accessible route for newly emerged microproteins to enter existing cellular networks.

## Introduction

Microproteins constitute an increasingly recognized class of proteins, typically fewer than 100 amino acids in length, translated from small open reading frames (sORFs), also termed translons^1^. Historically, sORFs were overlooked by conventional protein-coding gene annotation methods and they have remained largely uncharacterized due to their small size, presence of non-AUG start codons, overlap with canonical ORFs, and lack of sequence conservation and protein-level evidence^2^. Advances in ribosome profiling^3,4^, proteomics^5,6^ and comparative analyses of increasingly large collections of closely related, high-quality genomes have led to the identification of hundreds to thousands of putative sORFs across the tree of life^7–18^. sORFs have been identified in supposed untranslated regions of mRNAs, including upstream (uORF), downstream (dORF), internal to or overlapping with canonical coding sequences, or in long non-coding RNAs (lncRNAs) and intergenic regions. Comparative genomics^19^ and improved genome annotations^20^ have further revealed the potential of protein-coding genes arising *de novo* from previously non-coding DNA^21^. This has expanded our knowledge of the protein-coding potential of the genome and reshaped the view of how proteomes evolve. While short *de novo* genes are widespread in eukaryotic genomes^22^, most of them are rapidly lost and only a subset persists across millions of years of evolution and may acquire or refine biological function^21^. Emerging results suggest that some human-specific *de novo* sORF-encoded microproteins influence cellular phenotypes and play a role in disease^17,23,24^. However, whether newly emerged microproteins possess intrinsic molecular properties that enable them to engage in existing protein interaction networks, and how such interactions might contribute to the evolution of biological function, remains poorly understood.

A fundamental question is whether newly evolved microproteins adopt stable folded structures or exist primarily as intrinsically disordered proteins (IDPs). While some studies have suggested that microproteins are intrinsically disordered^25–27^, others have proposed that they can adopt folded structures^28–31^. Unlike evolutionarily established proteins, newly emerged sequences have had limited time to evolve sequence features required for stable, cooperative folding, potentially favouring disordered states during early protein evolution. Consistent with this idea, high-throughput FRET-FACS screening of putative human *de novo* microproteins conducted by Aubel et al.^32^ revealed that older microproteins show greater compactness than younger microproteins. These observations raise the possibility that intrinsic disorder is a common feature of young microproteins. If and how such newly emerged, predominantly disordered microproteins establish interactions with and gain access to existing cellular interaction networks remains largely unexplored.

Short Linear Motif (SLiM)-based interactions provide a potential mechanism by which intrinsically disordered microproteins may interact with cellular proteins. SLiMs are compact sequence patterns (3-10 aa stretches) that bind to folded peptide-recognition domains, often with low-to-mid micromolar affinities^33,34^. Their limited amino acid footprints allow for rapid evolutionary emergence^35,36^, offering an attractive potential route by which newly emerged microproteins may engage in pre-existing interaction networks. There is a widespread occurrence of SLiM-mediated interactions in the human interactome, with tens to hundreds of thousands of motif instances and thousands of SLiM-binding domains^33,34,37^. It is plausible that newly evolved microproteins contain SLiMs that contribute to the motif-based part of the human interactome. However, direct experimental evidence of SLiM-mediated interactions involving young microproteins remains limited, with only a handful of studies published to date^10,38,39^.

Here, we address these questions by combining biophysical characterization of the conformational properties of a select set of putative human *de novo* microproteins with a systematic analysis of their potential SLiM-based interactions. Using this integrated approach, we find that the examined microproteins are predominantly intrinsically disordered, with some propensities to take α-helical folds. We show that SLiM-like sequences are common in both *de novo* microproteins and their randomized sequence counterparts. Importantly, experimental analysis demonstrates that a subset of microprotein-derived motifs binds established peptide-binding domains with affinities within the range of canonical SLiM-mediated interactions. These findings provide direct support for a model in which newly evolved microproteins may acquire molecular function through disorder-enabled, motif-driven interactions, offering a framework for understanding how young microproteins may become functionally incorporated into existing cellular protein-protein interaction networks.

## Results

### Selection of *de novo* microproteins for experimental characterization

We based our selection of *de novo* microproteins on a previous study by Aubel et al.^32^, where the end-to-end compactness of human *de novo* microproteins retrieved from the sORFs.org database^8,40^ was assessed. We focused on the 30 microproteins with the highest apparent compactness in their screen (MP1-30) (**Figure 1a**, **Table S1.a**). The corresponding sORFs represent several genomic and transcript contexts, with most originating from uORFs, dORFs and lncRNAs, but some are derived from intronic and intergenic sequences (**Figure 1b**). Amongst the 30 sORFs included, 57% (17 out of 30) have been annotated as protein-coding in the sORFs.org database^8,40^. Independently of this annotation, 17 sequences (not necessarily the same ones) have a canonical start codon, and the complete set uses ten different start codons (**Table S1.a**). The evolutionary ages of the studied microproteins were inferred based on sequence homology in outgroup genomes. The largest fraction (33%) could be traced back to new world monkeys (**Figure 1c-d**), while five microproteins are human-specific. MP15 exhibited the deepest detectable conservation, with homologous sequences identified in eutheria. The selected microproteins range from 35 to 60 amino acids in length (**Table S1.a**), reflecting the size constraints of the original library used by Aubel et al.^32^.

**Figure 1.**
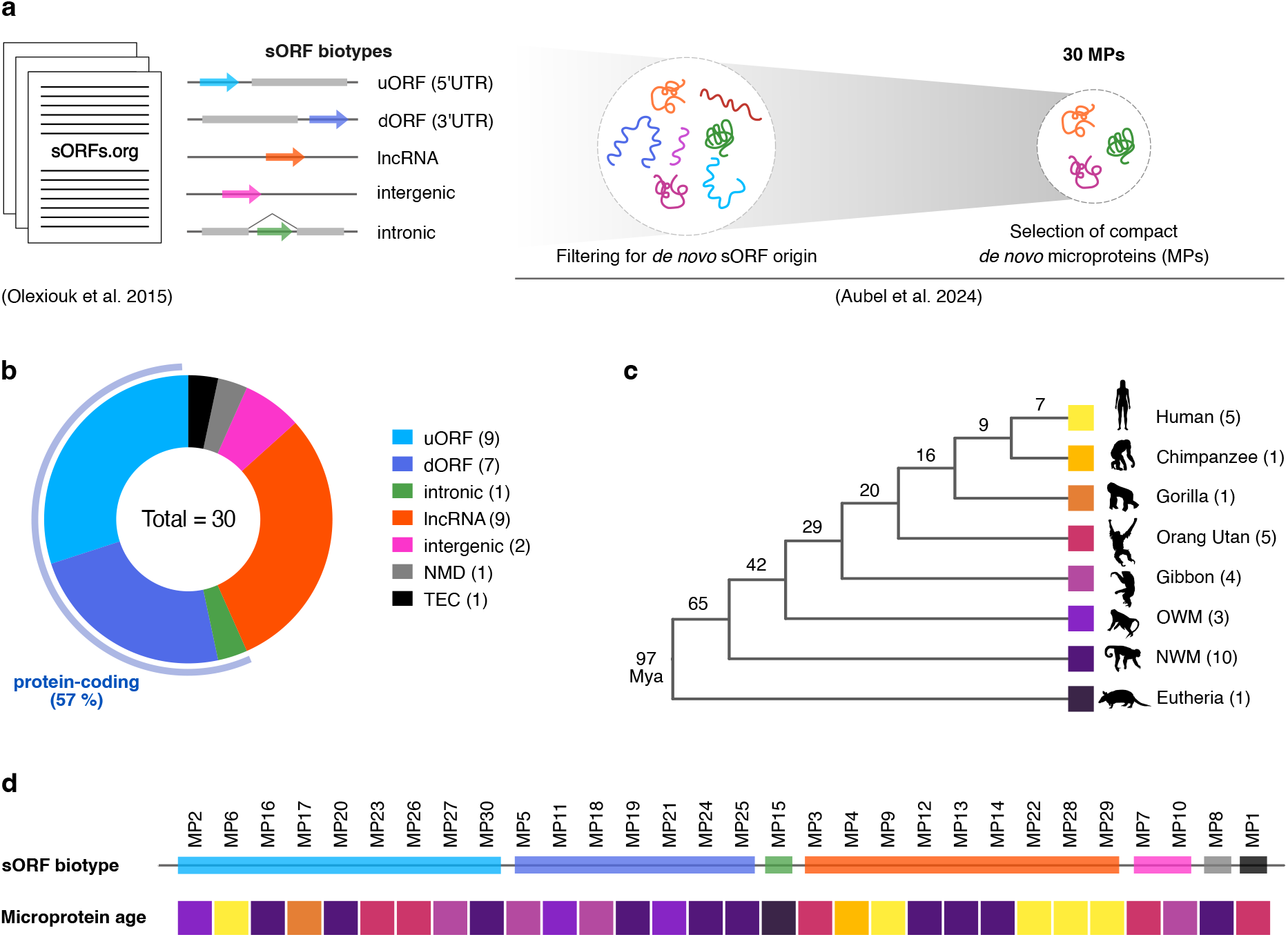
Characteristics of the selected *de novo* sORFs and their encoded microproteins. (**a**) Selection of the 30 most compact *de novo* small open reading frame (sORF)-encoded human microproteins from the Aubel et al study^32^. (**b**) Donut chart depicting the distribution of biotypes for the 30 *de novo* selected sORFs. The numbers in brackets indicate the number of sORFs of each biotype. 57% of the sORFs are annotated as protein-coding. NMD (=nonsense mediated decay); TEC (=to be experimentally confirmed). (**c**) Cladogram depicting the evolutionary relationship of taxa where microproteins were found to be conserved. The time of divergence for each species is indicated on branches (Mya=million years ago). Branch lengths do not reflect the evolutionary distance of the individual branches. The numbers in brackets indicate the number of microproteins conserved per species. (**d**) Transcript biotypes of the 30 *de novo* sORFs and the inferred evolutionary ages of their encoded microproteins, based on the phylogenetic analysis shown in **c**.

### *De novo* microproteins are predominantly disordered

At first, we explored the structural potential of the selected 30 microproteins. We predicted the structures with AlphaFold2 (AF2)^41^ (**Figure 2a, Figure S1**) as well as ESMFold^42^ (**Figure S2**). The predictions were dominated by low-confidence, non-globular conformations and consistent with a high degree of intrinsic disorder, although several sequences, such as MP14 and MP15, contained predicted α-helical regions with high confidence (pLDDT > 70), which agree with predictions of helicity by Agadir^43^ (**Figure S3**). The ESMFold predictions agreed with AF2 models and sequence-based predictions on overall predicted disorder as well as α-helical regions, where the latter tend to be predicted with lower confidence scores by ESMFold (**Figure S3**).

**Figure 2.**
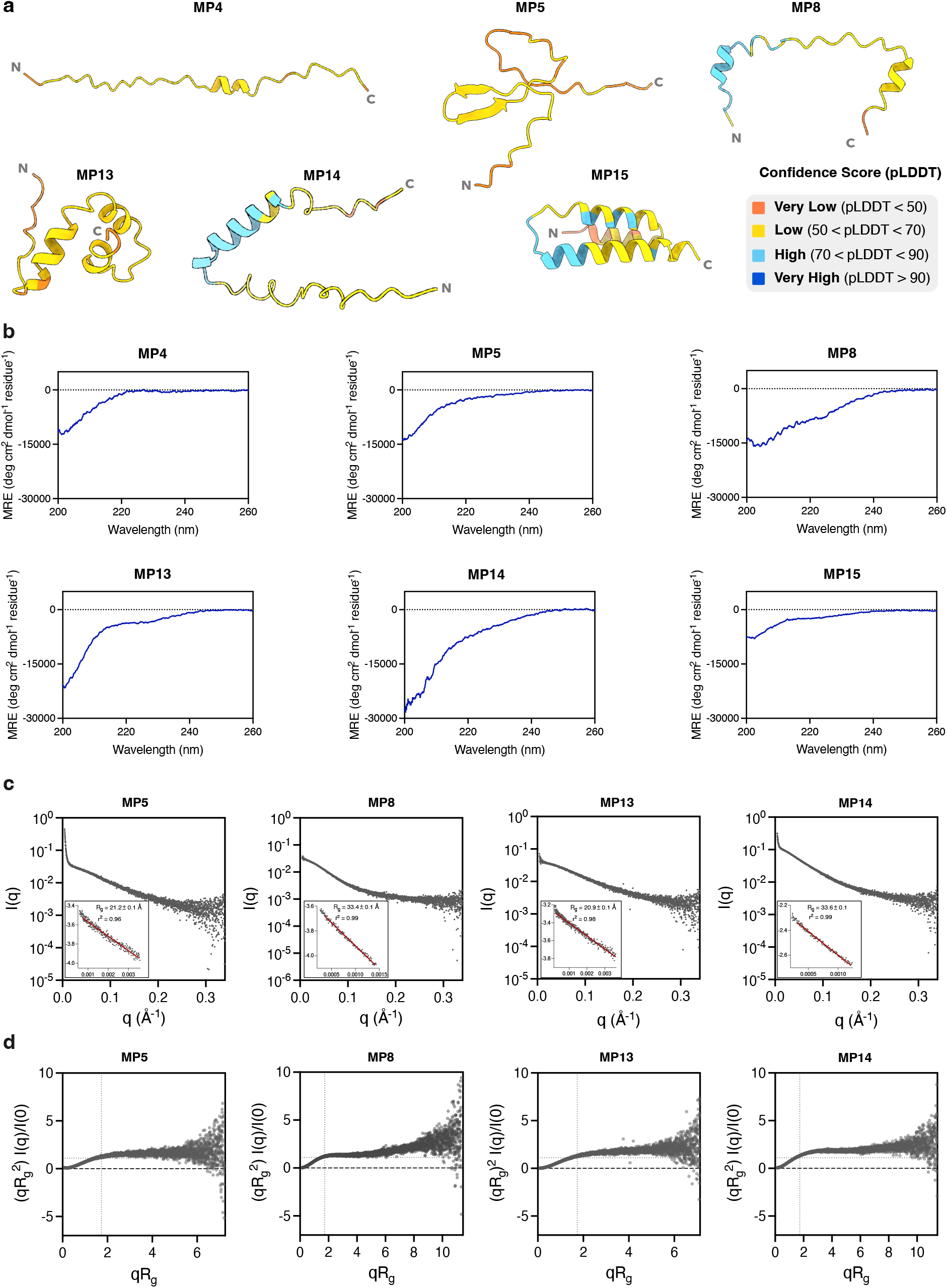
*De novo* microproteins predominantly adopt intrinsically disordered conformations. (**a**) AlphaFold2 (AF2)-predicted structures of the six investigated *de novo* microproteins, coloured according to pLDDT scores^41^.(**b**) Circular dichroism (CD) spectra of the microproteins suggest they are intrinsically disordered. Data are averaged across 4 accumulations and shown as mean residue ellipticity (MRE), corrected for microprotein concentration and size, to enable comparison. (**c**) Experimental small-angle X-ray scattering (SAXS) curves for the 4 analysed *de novo* microproteins. Insets show the Guinier fit region with the corresponding radii of gyration (R_g_). (**d**) Dimensionless Kratky plots of four microproteins. Grey dashed lines indicate the maximum peak height observed for bovine serum albumin (BSA), included as a globular protein reference.

To experimentally characterize the structural properties, we obtained synthetic genes encoding the microproteins in fusion with a 6xHis-GST tag. While most of the proteins were successfully expressed and purified as fusion proteins, only a subset (6 out of 30; **Figure 2a**) were obtained in sufficient yields and quality for structural characterization after cleavage of the His-GST tag, a problem commonly encountered during recombinant production of *de novo*-emerged proteins^44^. We therefore focused our experimental structural analysis on this subset and assessed secondary structure content using circular dichroism (CD) spectroscopy. All tested microproteins produced spectra consistent with a high degree of intrinsic disorder, characterized by maximum negative mean residue ellipticity (MRE) around 200 nm (198-203 nm for all microproteins) and lack of a double peak at 208/222 nm characteristic for α-helices (**Figure 2b**). Nevertheless, the spectra are indicative of varying degrees of disorder. While the CD spectra for MP4-MP5 and MP15 showed low MRE at 222 nm (-218,-2527 and-2219 MRE, respectively) indicative of a lack of α-helices, MP8 and MP13-14 produced stronger signals at 222 nm (-7899,-3678 and-7023 MRE, respectively) in the spectra consistent with partial or transient α-helices. Thus, although the experimentally amenable microproteins are predominantly disordered, they differ in their sequence-encoded propensity to populate secondary structure.

To gain further insight into their structural properties, we performed small-angle X-ray scattering (SAXS) characterization of five microproteins (MP5, MP8, MP13 and MP14, **Figure 2c, d**; MP15, **Figure S4e**). For all five microproteins, data were collected over a concentration range of 0.2–2.0 mg/mL (**Table 1**). For MP5, MP8, MP13 and MP14, the shape of the scattering profiles showed no concentration dependence (**Figure S4a-d)**, and further analyses were done with the highest-concentration data. However, the raw scattering curves (**Figure 2c**, **Figure S4a-d**) show a slight upturn at low q values, which can indicate interparticle interactions^45^. Because of strong concentration effects (**Figure S4e**), the MP15 data was not analysable.

**Table 1.** SAXS data acquisition and experimental results for four *de novo* microproteins.

| **Acquisition Parameters** |  |  |  |  |
|---|---|---|---|---|
| Beamline | B21, Diamond Light Source, Oxfordshire (UK) |  |  |  |
| Detector | Eiger 4M (Dectris) |  |  |  |
| Beam size (mm) | 0.112 x 0.24 (horizontal x vertical) |  |  |  |
| Energy | 14 keV |  |  |  |
| Sample to detector distance (mm) | 3700 |  |  |  |
| q range (Å<sup>-1</sup>) | 0.0045-0.34 |  |  |  |
| No. acquisitions per sample | 21 |  |  |  |
| Temperature (K) | 293 |  |  |  |
| **Experimental Results** |  |  |  |  |
|  | **MP5** | **MP8** | **MP13** | **MP14** |
| Concentration (mg/mL) | 2.0 | 1.5 | 2.0 | 2.0 |
| R<sub>g</sub> [from Guinier approximation] (Å) | 21.2 ± 0.1 | 33.4 ± 0.1 | 20.9 ± 0.1 | 33.6 ± 0.1 |
| qR<sub>g</sub> limits [from Guinier approximation] | 0.57-1.26 | 0.63-1.25 | 0.35-1.25 | 0.59-1.17 |
| r<sup>2</sup> [from Guinier approximation] | 0.96 | 0.99 | 0.98 | 0.99 |
| R<sub>g</sub> [from calculated scattering curve] (Å) | 21.9 | / | 19.6 | / |
| R<sub>g</sub> [from P(r)] (Å) | 30.4 ± 2.6 | 36 ± 0.3 | 23.4 ± 0.5 | 38.5 ± 0.7 |
| Molecular mass (kDa) |  |  |  |  |
| from amino acid sequence | 6.3 | 5.1 | 6.3 | 6.1 |
| from BioXTAS-RAW (Bayes) | 8.5 (6.7-9.1) | 19.9 (19.6-23.4) | 7.3 (6.7-9.1) | 18.7 (17.8-20.9) |
| From SAXS-MoW | 6.3 | 30.6 | 4.8 | 24.0 |

The four analysed microproteins show large radius of gyration (Rg) values (**Figure 2c**, **Table 1**). Together with the plateau-shaped dimensionless Kratky plots (**Figure 2d**) this indicates that the microproteins are disordered in solution, consistent with the CD data. The R_g_ values estimated from the Guinier fit and the pair distance distribution (P(r)) were generally in good agreement, supporting sample quality^46^, and consistent with a flexible system (**Table 1**, **Figure S4a-d**). Additionally, we estimated the molecular mass of the microproteins (**Table 1**). For MP5 and MP13, the experimentally estimated masses were close to those calculated from their amino acid sequences. Comparison of calculated scattering curves for the predicted monomeric structures of MP5 and MP13 to the experimental SAXS data shows that both protein models can explain the data when regions lacking secondary structures are allowed flexibility (**Figure S4f, g**), indicating that the degree of structure in these models is consistent with their solution structures. In contrast, the apparent molecular masses of MP8 and MP14 were 3-to 6-fold higher than expected (**Table 1**), potentially arising from oligomerization or aggregation.

Taken together, even though the *de novo* microproteins were selected for their high apparent compactness^32^, the CD and SAXS measurements indicate that the experimentally accessible *de novo* microproteins are predominantly disordered, although some exhibit a propensity to populate α-helical conformations and others show tendency for oligomerization or aggregation.

### *De novo* microproteins display distinct intrinsic helical propensities and show distinct TFE-induced transitions

We next sought to characterize the intrinsic folding propensity of the microproteins in greater detail. Because neither the computational predictions nor the experimental data indicated substantial β-strand content, we focused on α-helical structures. To this end, we used 2,2,2-trifluoroethanol (TFE) as a probe for the intrinsic helical propensity^47–50^. TFE serves as a cosolvent, or nanocrowding agent, that stabilizes local α-helical conformations, providing a means to probe the latent helical propensity of IDPs^51^. The microproteins displayed different behaviours in the presence of TFE (**Figure 3a**, **Figure S5a**). MP4 and MP5 showed no or little gain in α-helical signal across the TFE titration, whereas MP8, and MP13-15 displayed progressively stronger increases in helicity as indicated by enhanced negative ellipticity at 208 and 222 nm. Thus, there are substantial differences in the capacity of the microproteins to adopt α-helical conformations under helix-promoting conditions.

**Figure 3.**
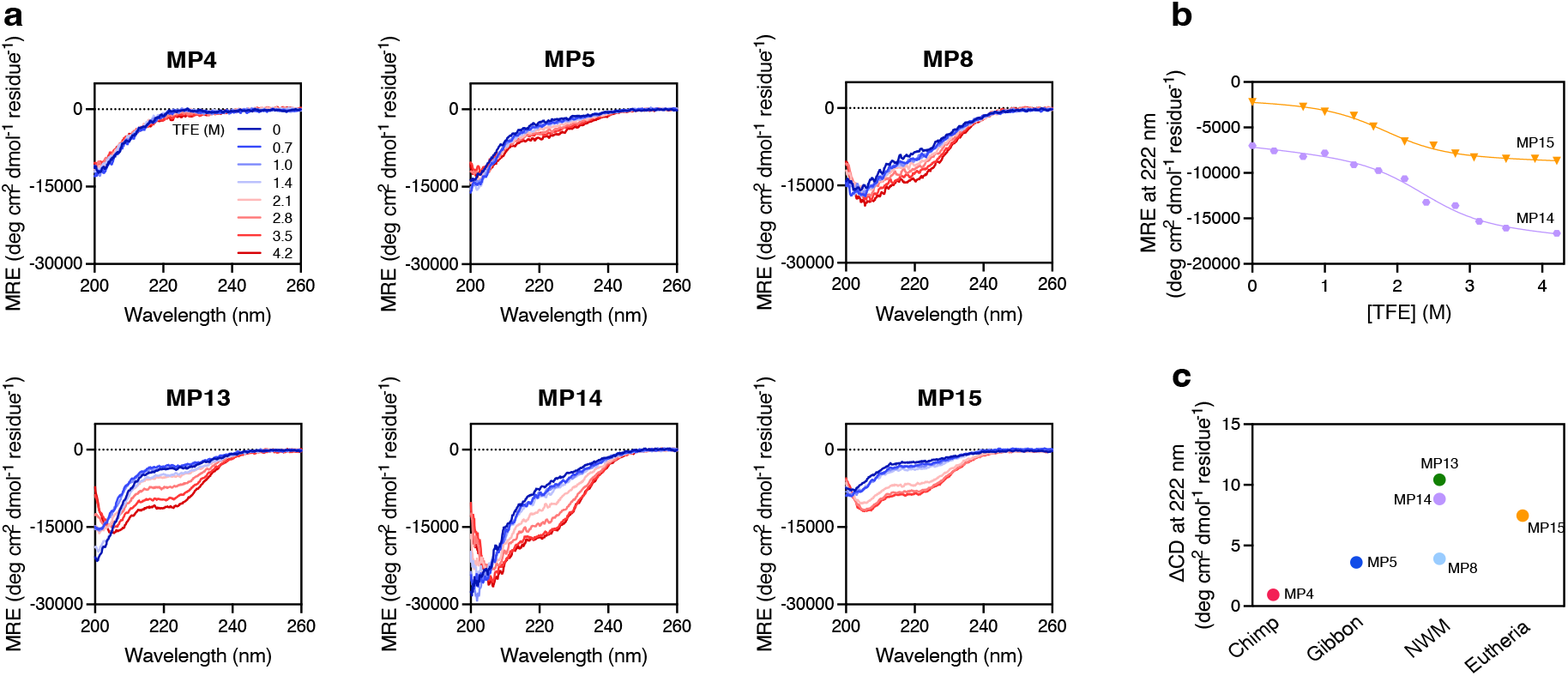
*De novo* microproteins have varying intrinsic helical propensities and show distinct cooperative and non-cooperative TFE-induced folding transitions. (a) Circular dichroism (CD) spectra of six *de novo* microproteins acquired at increasing 2,2,2-trifuloroethanol (TFE) concentrations (0 M TFE, dark blue; 4.2 M TFE, dark red). Data are averaged across four accumulations and shown as mean residue ellipticity (MRE) to enable comparison. (b) Cooperative folding transition of MP14 and MP15 upon addition of TFE. The data were fitted to a two-state folding model to determine the m-value (m_F-U_) and midpoint ([TFE]_50%_) of the folding transition. (c) Relationship between helical propensity (expressed as gain in CD signal at 222 nm from 0 M to 4.2 M TFE) and the inferred evolutionary age of *de novo* microproteins.

For IDPs, gradual non-cooperative transitions are expected when individual helical segments are stabilized independently rather than through formation of a concerted tertiary structure^51,52^. To examine these transitions in more detail, we monitored the CD signal at 222 nm across the TFE concentration series. TFE-induced helix formation can occur either gradually or through a more cooperative transition^53^. MP5, MP8 and MP13 exhibited non-cooperative behaviour, as indicated by a lack of a single isodichroic point in the CD spectra (**Figure 3a**) and non-sigmoidal titration curves (**Figure S5b**). These proteins likely populate multiple conformational states as their helical content increases with TFE concentration. On the other hand, the data for MP14 and MP15 were consistent with more cooperative structural transitions, as evidenced by a sigmoidal titration curve (**Figure 3b**) and the presence of a single isodichroic point in their CD spectra (**Figure 3a**). We fitted these data to a two-state folding model^53^, whereby the native unfolded (*U_Native_*) and the TFE-folded (*F_TFE_*) are the dominant populated states. From the fitting, we calculated the m-value, which reflects the change in solvent-accessible surface area and is expected to be larger for proteins that fold into more compact structures, and the midpoint of the transition ([TFE]_50%_), which represents the TFE concentration required to drive the coil-to-helix transition halfway, and is inversely proportional to a protein’s helical propensity. The m-values obtained for MP14 and MP15 (1500 ± 260 and 2900 ± 1100 cal mol^-1^ M^-1^, respectively) are typical for small cooperatively folding globular proteins^53,54^. Along with the midpoint values (2.3 ± 0.1 and 2.4 ± 0.1 M, respectively) they suggest that these sequences can undergo coordinated structural rearrangements rather than only independent local helix formation in the presence of TFE.

We next explored whether TFE-induced helical propensity was associated with the inferred evolutionary ages of the microproteins. Within the limited set of studied microproteins examined, the younger proteins (e. g. MP4 and MP5) show no or little gain in helical content upon TFE titration, whereas the older microproteins MP13-15 displayed greater TFE-induced helical content (**Figure 3c**). Although the sample size precludes firm conclusions about evolutionary trends, these observations are consistent with the possibility that secondary structure propensity may increase during the evolution of some *de novo* proteins.

Together, the results show that predominantly disordered *de novo* microproteins have distinct intrinsic α-helical propensities, which may potentially be stabilized by specific molecular environments or binding events.

### *Both de novo* microproteins and random sequences harbour putative SLiMs

Having characterized the structural properties, we next investigated the potential of the studied microproteins to engage in interactions with established SLiM-binding domains. To generate an experimentally testable set of predicted interactions we employed our available large-scale information on SLiM-binding determinants of 217 human protein domains from 34 domain clans^37,55^. We scanned the 30 *de novo* microprotein sequences for matches to the domains’ specificity determinants (**Figure 4a**, **Table S2**). The analysis suggested candidate SLiMs in 29 out of 30 *de novo* microproteins, corresponding to over 500 (569) potential interactions with 126 human domains from 26 domain clans (**Figure 4a** and **b**, **Table S2**). Notably, MP15, which is the oldest microprotein in the set, did not contain any putative motif matches. Predicted interactions included domains belonging to several established peptide-recognition families, including the WW, SH3 (Src homology) and PDZ (PSD95/Dig1/ZO-1) domain families and members of the ubiquitin clan (**Figure 4b**). Because short and degenerate sequence motifs can arise readily by chance^35,56^, we performed the same motif-scanning analysis on 30 randomized sequences from the Aubel et al. study^32^ (**Table S1.b)**, matched to the *de novo* microproteins in length and amino acid composition (**Table S2**). Similarly to the search of the *de novo* sequences, putative motifs were found in all 30 random sequences suggesting just over 400 (433) interactions with 120 human domains from 27 domain clans (**Figure S6a**), thus slightly fewer than in the *de novo* microproteins (**Figure S6b**). To assess the statistical significance of individual motif occurrences, we performed a permutation test using 1,000,000 shuffled sequences preserving sequence length and amino acid composition. Only one MP motif instance, a putative MDM2 E3 ligase SWIB domain-binding motif in MP4, showed a nominal deviation relative to the random expectation (p = 0.006) (**Table S2.e**).

**Figure 4.**
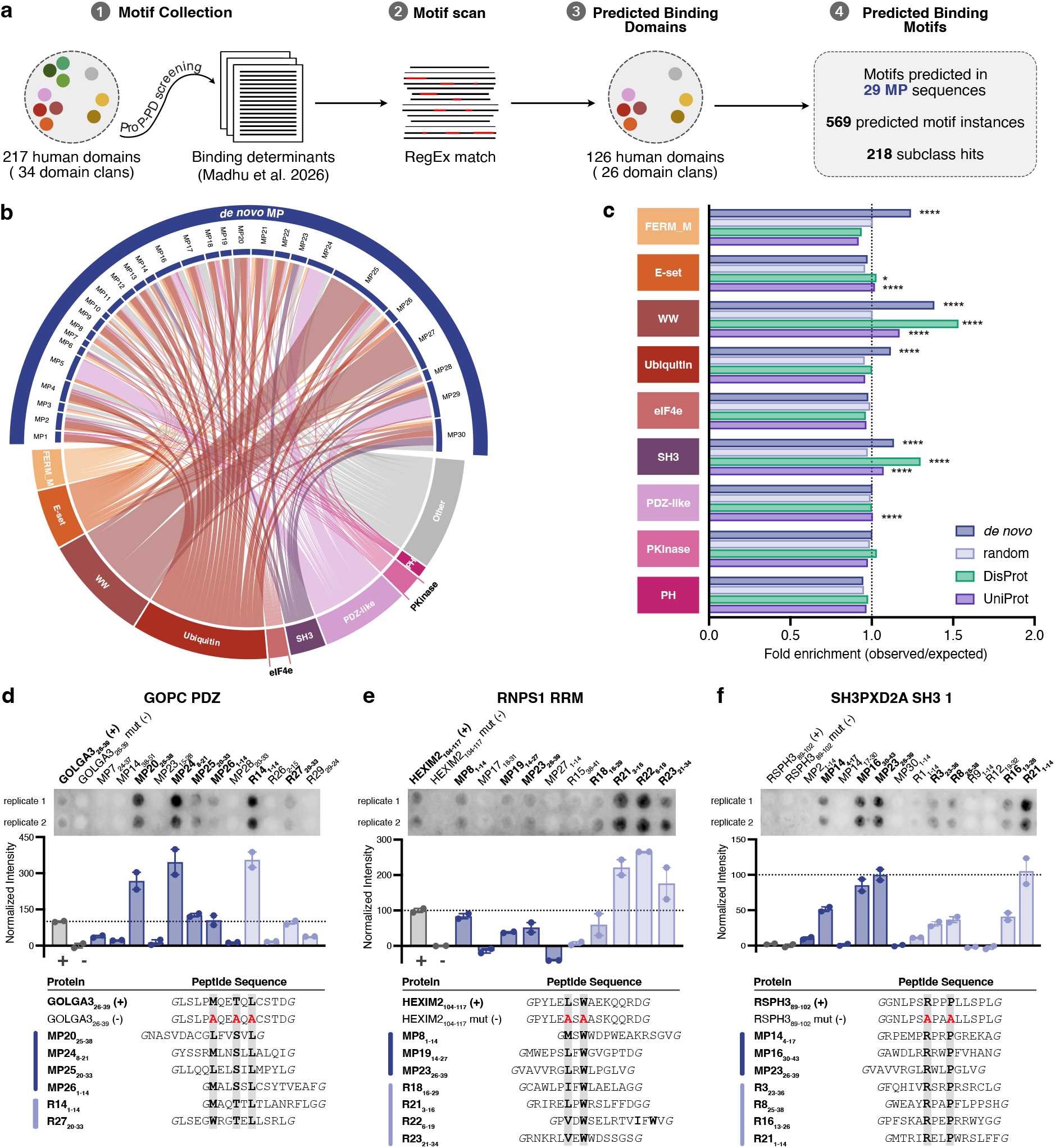
*De novo* microproteins frequently contain SLiM-matching sequences, a subset of which mediate peptide binding. (**a**) Schematic representation of short linear motif (SLiM) scanning in *de novo* microprotein (MP) sequences. An in-house curated collection of regular expressions (RegEx) for 217 domains (based on ref Madhu et al^37^), representing 34 domain clans, was used as the reference for finding matching motifs in MP sequences, as well as their randomized (R) sequence counterparts. (**b**) Chord diagram illustrating all predicted interactions of *de novo* MP sequences with 28 domain clans. Individual chords represent unique MP – domain interaction pairs. The thickness of the chords for each clan reflects the total number of unique MP – domain interactions, per clan. (**c**) Enrichment of SLiM-matching sequences across domain clans in *de novo* microproteins, randomized controls, DisProt and UniProt sequences. Fold enrichment was calculated as the ratio of observed to expected motif occurrences, with expected counts derived from a first-order Markov model fitted independently to each sequence set. Overlapping matches to different RegEx within the same clan were counted as a single motif occurrence when starting at the same position. The dashed line marks fold enrichment of 1. Stars: one-sided Poisson test, Benjamini-Hochberg correction over the 34 clans within each set (* q <= 0.05, ** q <= 0.01, *** q <= 0.001, **** q <= 0.0001). Stars mark enrichment only; no star means not significantly enriched. Owing to its large size (510,643 sequences), the UniProt dataset yielded statistically significant enrichment even for small effect sizes (e.g., E-set, 1.02-fold). (**d-f**) Results of the peptide SPOT array analysis for the (**d**) GOPC PDZ, (**e**) RNPS1 RRM, and (**f**) SH3PXD2A SH3 1 domains. Intensities were quantified and normalized against the positive and negative controls, except for SH3PXD2A where the controls failed (data were normalized against strongest and weakest MP binder). Data are represented as mean ± range of 2 replicates. Positive (+) and negative (-) controls are indicated. Binding peptides are highlighted in bold. Sequences of binding peptides are given below (MP, dark blue; R, light blue), with the binding determinants highlighted in bold.

We further expanded the motif-analysis to the complete set of 3,750 *de novo* microproteins and 3,750 randomized sequences from Aubel et al.^32^, and for comparison also to 13,347 defined disordered regions from DisProt^57^ and to 510,643 proteins listed in UniProt with little predicted disorder. The observed number of matches were compared with the counts expected under a first-order Markov model fitted to each sequence set, which reproduces its amino acid composition and the preferences between neighbouring residues. Most motifs occurred about as often as expected. However, 26 of the 106 motifs were enriched in *de novo* sequences (q ≤ 0.05), compared with 1 in random sequences, 54 in DisProt regions and 66 in UniProt proteins, and fold enrichments were higher in *de novo* than in random sequences (paired Wilcoxon signed-rank test across the 176 motifs, P = 5 × 10⁻^4^) (**Figure S6c**). Analysis by peptide-binding domain clans revealed that *de novo* microproteins were particularly enriched in putative WW-binding motifs (1.38-fold). DisProt regions similarly showed enrichment of WW-binding (1.5-fold) and SH3-binding motifs (1.37-fold). In contrast, motif frequencies in UniProt proteins generally remained close to expectation (0.92–1.17-fold) (**Figure 4c**).

Together, these analyses show that SLiM-like sequences are widespread in both *de novo* microproteins and randomized controls. The broader sequence-level analysis reveals an enrichment of specific motif classes in *de novo* microproteins that is not observed to the same extent in composition-matched randomized sequences. Whether these predicted motifs are capable of binding peptide-binding domains, however, cannot be inferred from sequence matches alone, but requires experimental testing.

### Validation of binding SLiMs in both *de novo* microproteins and random sequences

To test whether the predicted SLiMs are capable of binding their putative interaction partners, we performed peptide SPOT array experiments. We focused on three domains for which multiple microprotein-derived binding peptides had been predicted: the PDZ domain of GOPC (Golgi-associated PDZ and coiled-coil motif-containing protein) (**Figure 4d**), the RRM domain of RNPS1 (RNA-binding protein with serine-rich domain 1) (**Figure 4e**), and the first SH3 domain of SH3PXD2A (SH3 and PX domain-containing protein 2A) (**Figure 4f**). For each domain, we tested 5–8 microprotein-derived peptides matching the corresponding consensus recognition motifs (**Table S2.a**), together with additional candidates identified using less stringent motif criteria. Previously characterized ligands and motif-disrupting variants were included as positive and negative controls, respectively. For each domain, 3-4 microprotein-derived peptides were confirmed as well as the positive controls for two of three cases (**Figure 4d** and **e**). Several microprotein-derived peptides produced signals comparable to or stronger than those of the reference ligands. We also tested binding of candidate motifs identified in the randomized sequences against the same three domains. This validated 2-5 interactions per domain. Notably, for two of the three domains, more binding peptides were confirmed among the randomized sequences than among the corresponding *de novo* microprotein sequences (**Figure 4e** and **f**), and several randomized peptides produced comparatively stronger SPOT array signals. Although SPOT array intensities provide only a semi-quantitative measure of binding, these results demonstrate that both *de novo* microproteins and randomized sequences can harbour motifs, highlighting the potential for SLiM-mediated interactions to emerge in sequences without an evolutionary history.

### Affinities of *de novo* microproteins SLiMs span a broad affinity range and are easily evolvable

We next determined the affinities of putative SLiM-based microprotein interactions with five distinct protein domains using a competitive fluorescence polarization (FP) displacement assay (**Table 2**). We included interactions validated through the peptide SPOT array experiments as well as additional interactions judged to be of interest due to the function of the target proteins (e.g. the NTF2 domain of the stress granule protein G3BP1, the Mu homology domain (MHD) of the clathrin adaptor subunit AP2M1, and the SWIB domain of the E3 ligase MDM2). The candidate ligands are modelled to bind to the canonical peptide-binding pockets on each domain based on AlphaFold3 (AF3)^58^ modelling (**Figure 5a-d**, **Figure S8**). Microprotein-derived peptide affinities were successfully determined for 4 out of 5 probed domains (**Table 2**), and were found in most cases to bind with low micromolar affinities, as expected for SLiM-mediated interactions^34^. In several cases, microprotein peptides bound to their respective target domains with affinities close to, or higher than reference peptides of the domains (a PEX14 peptide for AP2M1 MHD domain; and a HEXIM2 peptide for RNPS1 RRM domain)^37^. Interestingly, the MP19_13-28_ peptide binds to the RRM domain of RNPS1 with a ∼10-fold higher affinity than its reference binding peptide HEXIM2_103-118_ (**Figure 5b, Figure S8b**). Thus, some recently emerged microprotein sequences already contain peptide motifs capable of binding established human peptide-binding domains with affinities comparable to known ligands.

**Figure 5.**
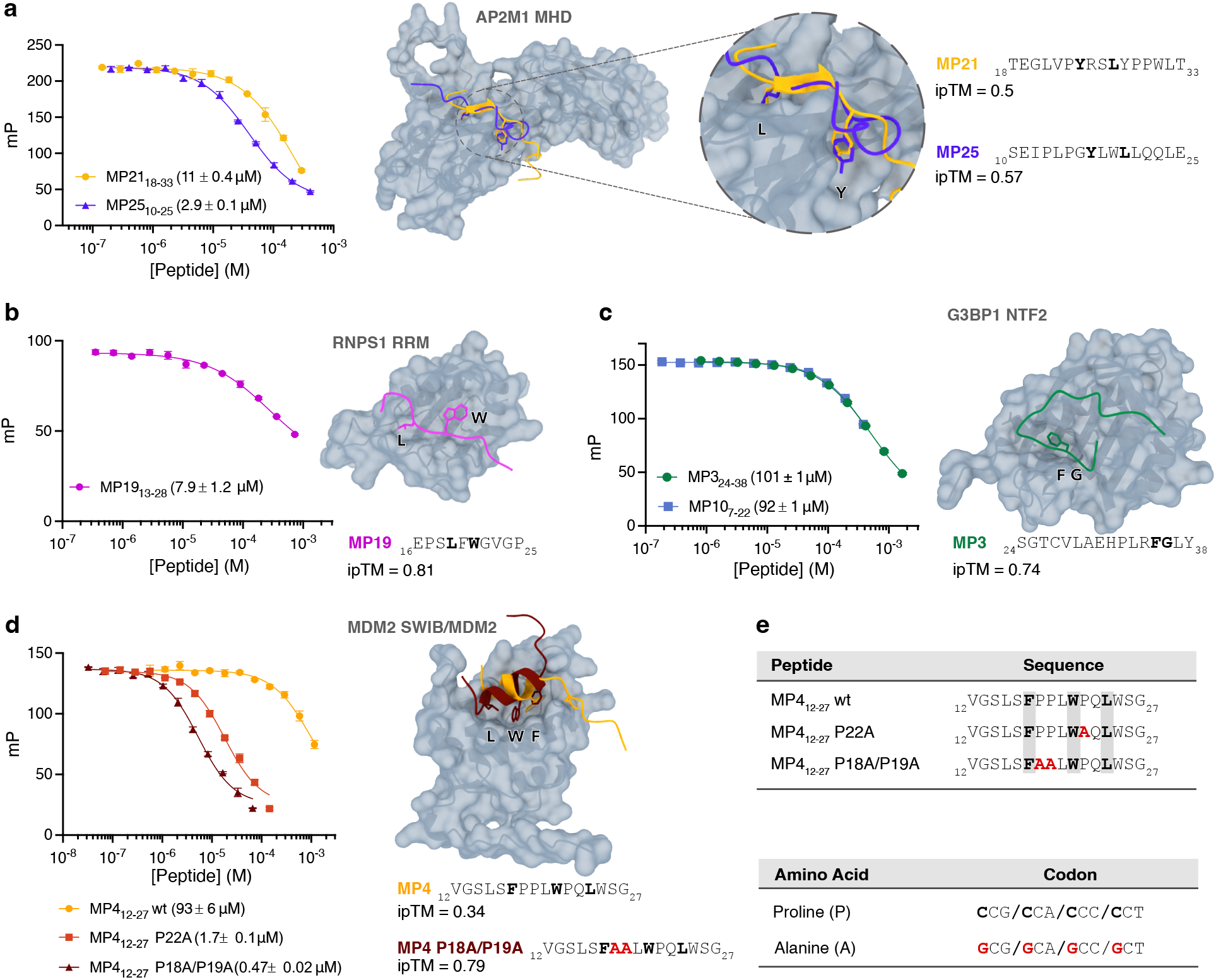
Affinities of *de novo* microprotein SLiMs span a broad affinity range and are easily evolvable. (**a-d**) Affinity measurements through competitive FP-based assay together with AlphaFold3 (AF3)-predicted structures of the target domains and their respective microprotein-derived peptide binders for (**a**) AP2M1 MHD, (**b**) RNPS1 RRM, (**c**) G3BP1 NTF2, and (**d**) MDM2 SWIM/MDM2 domains. FP data and the affinity values for the peptides are presented as mean ± SEM of 3 replicates. Predicted structures show confirmed peptide binders for each domain, and the key motif residues are shown. Sequences of the peptides used for the structural predictions and the ipTM confidence scores for the individual models are given. Key motif residues are highlighted in grey. (**e**) Amino acid sequences for wild-type (MP4_12-27_ wt), P22A mutant (MP4_12-27_ P22A) and P18A/P19A mutant (MP4_12-27_ P18A/P19A) peptides. A single nucleotide change, highlighted in red, is required for a proline-to-alanine mutation.

**Table 2.** Affinities of *de novo* microprotein SLiMs for their predicted domain binding partners as determined with fluorescence polarization (FP)-monitored binding experiments.

| **Bait Domain** | **Consensus Motif** | **Microprotein Peptide** | **Peptide Sequence** | **K<sub>d</sub> (µM)** |
|---|---|---|---|---|
| AP2M1 MHD | Y..L | MP21<sub>18-33</sub> | <sub>18</sub>TEGLVP**Y**RS**L**YPPWLT<sub>33</sub> | 11 ± 0.4 |
|  |  | MP25<sub>10-25</sub> | <sub>10</sub>SEIPLPG**Y**LW**L**LQQLE<sub>25</sub> | 2.9 ± 0.1 |
| RNPS1 RRM | [ILMV].W | MP8<sub>1-16</sub> | <sub>1</sub>**M**S**W**WDPWEAKRSGVLD<sub>16</sub> | N.B. |
|  |  | MP19<sub>13-28</sub> | <sub>13</sub>PMWEPS**L**F**W**GVGPTDV<sub>28</sub> | 7.9 ± 1.2 |
| G3BP1 NTF2 | FG | MP3<sub>24-38</sub> | <sub>24</sub>SGTCVLAEHPLR**FG**L*Y*<sub>38</sub> | 101 ± 1 |
|  |  | MP10<sub>7-22</sub> | <sub>7</sub>YSSPSREVTFWC**FG**EN<sub>22</sub> | 92 ± 1 |
| NEDD4 WW 2 | [LP]P.[Y] | MP25<sub>10-25</sub> | <sub>10</sub>SEIP**LP**G**Y**LWLLQQLE<sub>25</sub> | N.B. |
|  |  | MP27<sub>6-21</sub> | <sub>6</sub>PQGGSD**LP**Q**Y**RQGAWG<sub>21</sub> | N.B. |
| MDM2 SWIB/MDM2 | F...W..L | MP4<sub>12-27</sub> wt | <sub>12</sub>VGSLS**F**PPL**W**PQ**L**WSG<sub>27</sub> | 93 ± 6 |
|  |  | MP4<sub>12-27</sub> P22A | <sub>12</sub>VGSLS**F**PPL**W**AQ**L**WSG<sub>27</sub> | 1.7 ± 0.1 |
|  |  | MP4<sub>12-27</sub> P18A/P19A | <sub>12</sub>VGSLS**F**AAL**W**PQ**L**WSG<sub>27</sub> | 0.47 ± 0.02 |

Other probed interactions, such as those with the NTF2 domain of G3BP1, showed relatively weaker affinities. For example, the G3BP1 NTF2 domain is known to bind peptides with affinities ranging from low to mid micromolar (Kd = 1.1 μM for N-SARS-CoV-2^59^; K_d_ = 46 μM for the E3 ubiquitin ligase TRIM25^59^), and the affinities of the microprotein peptides MP3_24-38_ and MP10_7-22_ are markedly lower (**Figure 5c**, **Figure S8c**). However, the most pronounced difference was observed for the MDM2 SWIB interaction with MP4_12-27_, which has a ∼5000-fold lower affinity than the reference p53–MDM2 SWIB interaction^60^ (**Figure 5d**, **Figure S8e**). This observation is consistent with the expectation that newly emerged sequences have not undergone evolutionary affinity maturation for a biologically relevant target.

We next investigated whether such weak interactions could be strengthened through a small number of sequence substitutions. AF3-based modelling of the MDM2 SWIB–MP4_12-27_ complex suggested clashes of the peptide with the domain, and an incorrect placement of the key motif residues (**Figure 5d**, **Figure S8e**). Three proline residues interspersed within the MDM2 F…W..L binding motif likely restrict the formation of the α-helix required to position the key motif residues into the SWIB binding pocket^61^. We inquired whether we could artificially evolve a higher affinity of the MP4–MDM2 SWIB interaction by mutating the prolines in the MP4_12-27_ peptide to alanines, to promote helix formation. We generated two MP4_12-27_ peptide variants targeting different proline positions, a single P22A mutant and a double P18A/P19A mutant (**Figure 5d** and **e**). Notably, each amino acid substitution is accessible through a single nucleotide change (**Figure 5e**). Mutating P22A into an alanine was sufficient to increase the affinity ∼50-fold, whereas the P18A/P19A double mutation resulted in ∼200-fold increase in affinity (**Table 2** and **Figure 5d**). AF3 structural predictions suggested that the MP4_12-27_ P18A/P19A mutant fits well into the MDM2 SWIB binding pocket (**Figure 5d**), and the key motif residues align perfectly with the canonical p53 binder (**Figure S8e**). Altogether, the data shows that the weak interaction between MP4 and MDM2 can be converted into a substantially stronger interaction, consistent with a high evolvability of microprotein SLiMs towards higher binding affinities.

In summary, the newly evolved microproteins studied here contain SLiMs that may be recognized by peptide-binding domains with affinities similar to canonical interactions. Lower affinity interactions may easily be evolved towards higher affinity, given a relevant selection pressure. Whether such interactions are realized in cells will additionally depend on factors including microprotein expression levels, localization, stability, accessibility of the motif, competition with endogenous ligands, and the cellular concentrations of the interacting partners.

## Discussion

Microproteins have gained increasing attention over the last decade, challenging our view of proteome evolution and highlighting the underestimated coding potential of the genome. Among these microproteins, some are newly evolved. The emergence of protein-coding sequences *de novo* poses a fundamental evolutionary problem: how can a newly translated sequence acquire molecular properties compatible with the existing cellular environment and, potentially, become functionally integrated into it? Here, we characterized a set of putative human *de novo* microproteins with respect to their structural properties and their capacity for motif-mediated interactions. Our results support a model in which the newly emerged microproteins are predominantly disordered and contain SLiMs capable of engaging in interactions with established peptide-binding domains. Because such SLiM-like sequences arise readily from the microproteins’ underlying sequence properties, they may provide an accessible starting point for the evolution of new interactions.

Despite being selected for tendency of compactness^32^, structural predictions of the 30 *de novo* microproteins investigated suggest that the studied microproteins are largely disordered. Consistent with these predictions, our experimental characterization of six *de novo* microproteins supports that they are disordered and adopt extended and flexible conformations in solution. Importantly, *de novo*-emerged proteins with higher disorder content and less propensity to aggregate may be easier to express and purify^44^, potentially introducing a bias for disordered microproteins in our characterized set. The same properties may be advantageous during the early stages of protein emergence because they can enhance solubility and reduce the formation of potentially toxic aggregates, thereby increasing cellular tolerability^25,32^. Acquiring a stable and compact fold may require longer evolutionary time and selection pressure, to which young *de novo* microproteins have not been exposed^62^. The small size of the microproteins investigated here may further provide an additional constraint on stable folding. Nevertheless, *de novo* microproteins are not necessarily restricted to disordered states, as several experimentally-characterized examples have been shown to form secondary structures and even stable hydrophobic cores^29–31,63,64^, illustrating the need for a case-by-case characterization. Consistent with this, the microproteins examined displayed markedly different responses to TFE stabilization, ranging from little detectable helical propensity to cooperative TFE-induced structural transitions. Within our limited dataset, the older microproteins tended to have higher α-helical propensity, in agreement with previous observations^32^. Still, our observation of varying helical propensities shows that, like conserved IDPs^65^, *de novo* disordered microproteins exist on a spectrum of intrinsic disorder and encode distinct latent propensity for folding, suggesting that specific molecular and cellular contexts may drive the formation and stabilization of secondary structures.

We further asked how the predominantly disordered microproteins might engage in cellular protein-protein interaction networks. A high degree of intrinsic disorder may provide *de novo* microproteins with opportunities to become integrated into canonical cellular networks and acquire function by encoding SLiMs that are accessible for recognition by peptide-binding domains^34^. Consistent with this, our bioinformatic search for SLiMs in the 30 *de novo* microprotein sequences identified putative motifs in all but one microprotein sequence. However, we also predicted putative motifs in their randomized sequence counterparts. Computational analysis shows that SLiM-like sequences in *de novo* microproteins occur about as often as expected from amino acid composition. The mere occurrence of a sequence matching a known interaction motif is therefore not in itself evidence of functional selection. Nevertheless, *de novo* microproteins differed somewhat from random sequences of similar compositions, with proline-rich ligands of SH3 and WW domains being more frequent than expected. Notably, the proline-rich motifs come from proline-rich sequences, which more likely reflect the non-coding sequences from which these microproteins arose than selection for binding. Also established disordered regions in DisProt^57^ showed enrichment of proline-rich WW and SH3 motif content. Peptide SPOT array and FP experiments confirmed that a subset of the candidate motifs bind established peptide-binding domains. Several microprotein-derived peptides bound their targets with low micromolar affinities, in some cases comparable to established ligands. For example, peptides derived from MP21 and MP25 bound the clathrin adaptor AP2M1 within the affinity range reported for endogenous and viral ligands. Whether such interactions occur in cells remains unknown, but they illustrate that newly emerged sequences can encode biochemically relevant binding motifs without a long history of coevolution with the target protein.

The comparison with randomized sequences provides a further perspective on the interaction landscape. Randomized peptides frequently bound the tested domains and, for some interactions, produced stronger SPOT array signals than the corresponding *de novo* microprotein peptides or reference ligands. Because SPOT intensities provide only a semi-quantitative measure of binding, these differences should not be interpreted directly as affinity differences. Nevertheless, together with previous observations that young *de novo* microproteins differ measurably from unevolved random sequences^25^, the data raise the possibility that newly emerged proteins are subject to purifying selection against interactions that are excessively strong or otherwise deleterious. Under such a model, motif-like sequences arise frequently, whereas their interaction strengths and surrounding sequence contexts are subsequently attenuated, retained or enhanced according to their cellular consequences. Our analysis of the interaction of MP4 with the E3 ubiquitin ligase MDM2 shows that single nucleotide mutations can result in orders of magnitude increase in affinities and demonstrates evolvability of microprotein SLiM-mediated interactions.

In summary, our study provides insight into structural and binding properties of human *de novo* microproteins. We propose a framework where putative SLiMs occur frequently by chance in intrinsically disordered *de novo* microproteins, and serve as an evolutionary reservoir of potential interactions, which may be selected for if beneficial, or purged if deleterious to the cell. In this framework, a high degree of disorder enables accessibility of SLiMs to engage in interactions with the conserved proteome, which may facilitate the integration of microproteins into cellular networks.

## Materials and Methods

### Prediction of Protein Properties

Intrinsic disorder of the *de novo* microprotein sequences was predicted using the online server IUPred3^66^. Helicity of the sequences was predicted locally using the pyagadir implementation of AGADIR^43^ in Python3. Three-dimensional structures of microproteins were predicted using the AF2 implementation on ColabFold v1.5.5^67^ on Jupyter^68^. The template used was pdb100 and the multiple sequence alignment was set to single sequence as *de novo* microproteins are not expected to have homologous sequences. All other parameters were set to their default settings. Similarly, ESMFold^42^ structure predictions were performed on Jupyter notebook^68^, using ESMFold v.1 as the prediction model. The AF3 online server was used for the modelling of complexes between protein domains and microprotein-derived peptides^58^. Analysis and visualization of predicted structures was done in ChimeraX-1.10^69^.

### Age Estimation of *De Novo* sORFs

To verify age estimations and identify non-coding homologous sequences for the sORFs, the presence of the ORF was manually examined using the UCSC Genome Browser (http://genome.ucsc.edu)^70^. For all sORFs, the 30 Mammals UCSC MultiZ genome alignment^71,72^ was used, for sORFs 20, 14 and 15 the UCSC 100 Vertebrates UCSC MultiZ genome alignment^71^ was used additionally to identify non-coding homologs. For all sORFs, presence of a stop codon was verified up to 10 nucleotides after the human stop codon. Presence of ATG was verified for sORFs starting with ATG; for sORFs starting with non-canonical start codons (TTG, CTG, GTG) any non-canonical start codon was accepted. Start codons were considered up to 10 nucleotides before the human start codon. For sORF14, the only sORF containing introns, only exon sequences were considered. Because of the short ORF length and experimental focus, only full length ORFs were considered and homologous sORFs with premature stop codons were discarded.

### Protein Production

Sequences encoding the 30 *de novo* microproteins and randomized sequences (**Table S1.a** and **S1.b**, respectively) were obtained as N-terminally 6xHis-GST-tagged fusion proteins in pETM33 plasmid with kanamycin resistance from GenScript. Microproteins were produced recombinantly in *E. coli* BL21(DE3) in 2xYT medium (5 mg/mL NaCl, 10 mg/mL yeast extract, 16 mg/mL peptone) supplemented with 50 μg/mL kanamycin at 37 °C. At OD_600_ = 0.6-0.8, microprotein expression was induced with 0.1-1 mM Isopropyl β-D-1-thiogalactopyranoside (IPTG). The microproteins were expressed at 18 °C for 16 h, and the cultures were harvested by centrifugation at 4,000 x g, 4 °C. Bacterial pellets were washed with 10 mL 1 x PBS buffer (8 mg/mL NaCl, 0.2 mg/mL KCl, 1.44 mg/mL Na_2_HPO_4_, 0.245 mg/mL KH_2_PO_4_, pH 7.4) per 1L culture and harvested at 4,000 x g, 4 °C.

Pellets were resuspended in lysis buffer (50 mM Tris, 300 mM NaCl, 5 mM DTT, 2 mM CaCl_2_, 4 mM MgCl_2_, 20 μg/mL DNase, 20 μg/mL RNase, 50 μg/mL lysozyme and cOmplete™ EDTA-free Protease Inhibitor Cocktail (Roche), pH 7.8) using a sonication probe for 2 cycles of 60 s sonication with 1 s pulses. The lysate was clarified with centrifugation at 20,000 x g, 4 °C for 1 hr and the soluble fraction was filtered through a 0.22 μm sterile filter and incubated with glutathione affinity resin (Pierce™ Glutathione Agarose, ThermoFischer Scientific). Proteins were purified on the glutathione affinity resin with Tris buffer (50 mM Tris, 300 mM NaCl, pH 7.8), and the 6xHis-GST tag was enzymatically removed on the resin using in-house purified HRV-3C protease at 4 °C overnight.

For structural analysis, cleaved microproteins were further purified with gel filtration chromatography on HiLoad™ 16/600 Superdex 75 pg (Cytiva Life Sciences), pre-equilibrated with 50 mM sodium phosphate buffer (10.107 mg/mL Na_2_HPO_4_ x 7H_2_O, 1.679 mg/mL NaH_2_PO_4_ x H_2_O, pH 7.4), at 1 mL/min. Fractions containing the pure microproteins were collected, supplemented with 1 mM Tris(2-carboxyethyl) phosphine (TCEP) for microproteins containing cysteines, and flash frozen in liquid nitrogen. Alternatively, microproteins were further purified by ion exchange chromatography on HiTrap™ Q anion exchange chromatography pre-packed columns (Cytiva Life Sciences), equilibrated with HEPES buffer (20 mM HEPES, 50 mM NaCl, pH 7.8) and eluted in a stepwise manner with HEPES buffer (20 mM HEPES, 100-500 mM NaCl, pH 7.8). Pure microproteins were dialyzed against 50 mM sodium phosphate buffer and flash frozen in liquid nitrogen.

### Circular Dichroism

CD measurements were performed on a JASCO J-1500 spectrophotometer using a quartz cuvette with a 1 mm pathlength (Hellma, Müllheim, Germany). All experiments were performed in 50 mM sodium phosphate buffer at 25 °C at roughly 20 μM microprotein concentration. For each microprotein, the exact concentration was determined immediately prior to measurements on the NanoDrop One^C^ spectrophotometer (ThermoFischer Scientific). Prior to microprotein measurements buffer spectra were acquired to check for residual signal. Microprotein spectra were recorded at 190/200-260 nm, with an interval of 0.1 nm, a digital integration time (D.I.T.) of 1 s, and a scanning speed of 50 nm/min. All spectra were averaged across 4 accumulations. Raw ellipticity data in millidegrees were converted to MRE using information on the size and concentration of the microproteins. Spectra for TFE titrations were acquired in the same manner. Prior to measurement, protein was mixed with TFE to obtain the required concentrations of TFE and incubated at room temperature for 10 min. All data was analysed and visualized in GraphPad Prism v.10.6.1. To obtain the TFE midpoint ([TFE]_50%_) and m-value (m_F-U_) of the cooperative folding transition, the data was fitted to a modified two-state folding model (**Equation 1**):

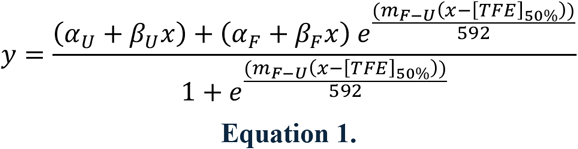

where *y* is the signal in mdeg and *x* is the TFE concentration. *α_U_* and *α_F_* represent the baseline at the unfolded (U) and folded (F) states, respectively, and *β_U_* and *β_F_* represent the slope of the baseline at the unfolded and folded states, respectively. Parameter *m_F_*_−*U*_ describes the slope of the cooperative folding transition. [*TFE*]_50%_ refers to the concentration of TFE at which 50% of the unfolded-to-folded transition has been completed and is known as the midpoint. The number 592 describes the term R*T at room temperature (298 K) in cal mol^-1^ K^-1^.

### Small Angle X-Ray Scattering

Purified microproteins were dialyzed against SAXS buffer (20 mM Tris, 150 mM NaCl, 1 % glycerol, pH 7.8, supplemented with 1 mM TCEP for microproteins containing cysteines) at 4 °C overnight, using the 1 kDa cut-off Mini dialysis kit (Cytiva, USA). To prepare the concentration series, the lowest concentration sample was prepared first before concentrating to the next higher concentration to avoid aggregates in the lower concentration samples. Microproteins were concentrated with the Vivaspin® 15R 2,000 MWCO centrifugal filters (Sartorius, UK) at 3,500 x g in 10 min increments. All samples were prepared using the dialysis buffer to ensure the best possible buffer match for data collection. Samples were flash frozen in liquid nitrogen.

Data was acquired on the B21 beamline at the Diamond Light Source Ltd synchrotron (UK)^73^. The data acquisition parameters are given in **Table 1**. Data was downloaded from the Diamond User Administration System, and all data processing was done in the BioXTAS RAW 2.4.1 software^74^. For each microprotein, scattering curves were recorded at multiple concentrations ranging from 0.2-2.0 mg/mL to check for concentration-dependent oligomerization or non-specific aggregation. Buffer scattering curves were recorded before and after acquisition of the microprotein scattering curves, averaged, and subtracted from the averaged microprotein scattering curves. Scattering curves at different concentrations were overlayed and if no concentration-dependent behavior was observed, we proceeded with analysis of the scattering data at the highest microprotein concentration. Outliers from the scattering curves were removed manually. Data for MP14 at 0.2 mg/mL and MP15 for 0.5 mg/mL were not processed because the raw scattering curves were of poor quality.

The R_g_ values were estimated from the experimental data using Guinier approximation. Experimental scattering curves were compared to calculated scattering curves derived from AF2-predicted monomeric structures of the microproteins using MultiFoXS^75^. Molecular mass of the microprotein samples was determined using a Bayesian inference method implemented in BioXTAS RAW as it has been shown to be a reliable concentration-independent method for molecular weight estimation^76^. Additionally, the molecular mass was also estimated using the SAXSMoW 2.0 online server^77^ and the results were compared to those of the BioXTAS RAW analysis. All plots relating to SAXS data were initially created in BioXTAS RAW and adjusted for publication in GraphPad v.10.6.1.

### Motif Scan

We curated a list of previously in-house generated information on motif recognition of peptide-binding domains (**Table S2.a**), published by Benz et al^55^. The motif collection is based on 217 human baits for which we were able to determine specificity determinants in ProP-PD selections^37^. To find matching motifs in microprotein sequences, we scanned the *de novo* and randomized counterpart sequences for sub-sequences matching the regular expression (RegEx) in our collection, allowing overlaps (**Table S2.b** and **c**). A permutation test against 1,000,000 shuffled sequences (preserving length and amino acid composition) was used to assess the probability of the RegEx matches, where two empirical p-values were calculated (**Table S2.e**), one for the fraction of shuffled sequences presenting equal or greater matches to the RegEx in the original sequence, and one for the presence of one or more matches (when the original sequence has only one match, these two p-values are the same).

### Statistical Analysis of Motif Occurrence

We tested whether the 106 unique RegEx of the 217 domains of the SLiM collection (34 domain clans) occur more often than expected in four sequence sets: the 3,750 *de novo* and the 3,750 length-matched random sequences of the library of Aubel et al.^32^, 13,347 disordered regions from DisProt^57^ (release 2025_06), and 510,643 UniProtKB/Swiss-Prot proteins (release 2025_04) after removal of DisProt entries and of proteins with more than 30% of residues predicted as disordered by IUPred3^66^. Observed occurrences, counting overlapping matches, were compared with the number expected under a first-order Markov model fitted to the amino acid and dipeptide frequencies of each set; expected counts were calculated exactly (SI Text). Enrichment was expressed as fold enrichment (observed/expected) and tested with a one-sided Poisson test, and P values were corrected across all motifs within each sequence set with the Benjamini–Hochberg procedure (q ≤ 0.05 considered significant). For the domain-level analysis (**Figure 4c**), all RegEx of a domain type were combined so that each site was counted once. As sensitivity analyses, we repeated the analysis with a composition-only (IID) model and with position-specific models (five relative position bins), and we compared *de novo* with random sequences directly by the number of sequences containing each motif (two-sided Fisher’s exact test). Enrichment profiles of the sequence sets were compared with paired Wilcoxon signed-rank tests. Equations and further details are given in the SI Text; the code is available at https://github.com/ArsLeicholt/slim_probability.

### SPOT Array

SPOT array membranes of predicted microprotein-derived binding peptides were purchased from JPT Peptide Technologies (Germany). To activate the membranes, they were incubated in methanol for 5 min and washed with TBST buffer (20 mM Tris, 150 mM NaCl pH 8.0, 0.05 % Tween-20) for 3 min, for three total washing rounds. Then, membranes were blocked in blocking buffer (TBST + 3 % BSA) for 1 h at 4 °C. The membranes were incubated with purified His-GST-tagged protein domains overnight at 4 °C, to allow for binding. Washing with TBST buffer was performed for 1 min, for a total of three washing rounds. HRP-conjugated anti-GST antibody (anti-GST HRP conjugatre, RPN1236, Cytiva) was added to the membranes for 1 h at 4 °C, after which the membranes were washed with TBST buffer for 3 x 1 min. A final wash with TBS buffer for 1 min was performed. Membranes were imaged using the ECL reagent (Clarity Max Western ECL substrate, 1705062, BioRad) on the ChemiDoc imaging system (BioRad). Images were processed using Fiji ImageJ v.1 software and all plots were prepared in GraphPad v.10.6.1.

### Fluorescence Polarization

FP affinity measurements were performed in black non-binding surface flat-bottom 96-well plates (Corning) on the SpectraMax iD5 microplate reader (Molecular Devices), with an excitation wavelength of 485 nm and an emission wavelength of 535 nm. All measurements were done in FP buffer (50 mM sodium phosphate buffer +/- 1 mM TCEP, 0.05 % Tween-20, pH 7.4), at 20-25 °C in technical triplicates, with a working volume of 50 μL/well. The experimental conditions were identical for all domains investigated, unless stated otherwise. Data analysis and plotting was done in GraphPad v.10.6.1.

First, the affinity of the investigated protein domains for their respective FITC-labelled peptide was determined in saturation binding experiments. The protein domain was serially diluted 1:1 with FP buffer and mixed with FITC-labelled peptide at 10 nM final concentration, such that [protein domain] >> [FITC-labelled peptide]. FITC-labelled peptide at 10 nM in 50 mM sodium phosphate buffer was used to determine the bottom baseline (*B_bottom_*). The G-factor was adjusted to obtain a value of 10-30 mP for the bottom baseline. Saturation binding curves from three technical triplicates were averaged and plotted. The dissociation constant (*K_d_*) was obtained by fitting the data to **Equation 2**:

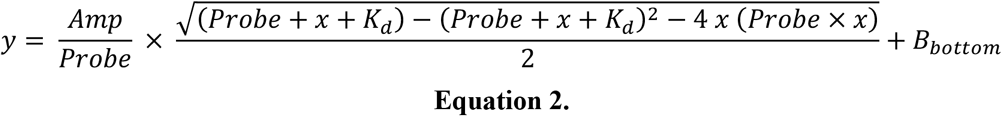

where *y* is the FP signal, *x* is the molar concentration of the investigated domain, *Probe* is the molar concentration of the FITC-labelled peptide, *Amp* is the amplitude of the FP signal (*B_top_* − *B_bottom_*), and *K_d_* is the dissociation constant.

Affinities of the investigated domains for the microprotein-derived peptides were determined in competitive displacement experiments of FITC-labelled probe peptides with unlabelled microprotein-derived peptides. In a similar manner as before, a serial dilution of unlabelled microprotein-derived peptides in FP buffer was prepared. To this, a pre-mixed solution of FITC-labelled peptide (10 nM) and domain of interest was added. The concentration of the domain in the mixture was set to be 2-10 x *K_d_* obtained in saturation binding experiments. Complex between FITC-labelled peptide and the domain of interest was used to determine the top baseline (*B_top_*). The displacement curves were plotted and fitted to a sigmoidal model (**Equation 3**) describing a competitive binding assay, adapted from Nikolovska-Coleska et al.^78^:

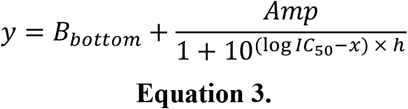

where *y* is the FP signal, *x* is the molar concentration of the unlabelled microprotein-derived peptide, ℎ is the hill slope of transition, and *IC*_50_ is the molar concentration of the unlabelled peptide at which 50 % of the FITC-labelled peptide is displaced from the domain of interest (equal to 50 % inhibition of the domain – labelled peptide complex).

From the resulting *IC*_50_ values from three replicates, three inhibitory constants (*K_i_*) for a given peptide were calculated based on the **Equation 4**, as described previously^78^:

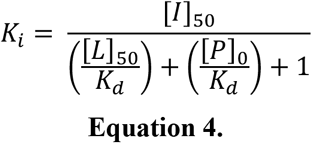

where *K_i_* is the inhibitory constant of the unlabelled peptide at 50 % inhibition, [*I*]_50_is the molar concentration of the free unlabelled peptide at 50 % inhibition, [*L*]_50_ is the molar concentration of the free labelled peptide at 50 % inhibition, [*P*]_0_is the molar concentration of the free protein domain at 0 % inhibition, and *K_d_* is the dissociation constant of the labelled peptide – domain interaction. The resulting *K_i_* values from three replicates were averaged to calculate the reported *K_i_* for a given microprotein-derived peptide–domain interaction. All *K_d_* and *K_i_* values are reported as the average of three replicates ± SEM.

## Acknowledgements

This work was supported by grants from the Swedish Research Council (Y.I.: 2024-04306; PJ: 2024-04884I). Y.I., E.B.B., and K.H. received funding from Volkswagen foundation grant code 98183. We are grateful to Diamond Light Source (Oxfordshire, UK) for access and time on beamline B21 under proposal MX37715.

## Author Contributions

I.M., P.J., E.B.B., K.H., and Y.I. conceived the study. I.M. produced proteins, performed CD and SAXS measurements and analysed the data. M.S. assisted with SAXS experiment planning and data interpretation. M.A. conducted age and *de novo* status estimation of the investigated microproteins. L.S. conducted computational motif scanning. L.E. carried out statistical analysis of motif occurrence. P.M. performed SPOT array experiments. I.M. and P.M. analysed SPOT array data. I.M. performed FP experiments and analysed the data. I.M. and Y.I. wrote the first draft. All authors reviewed and approved the manuscript. Y.I. coordinated the study.

## References

1. Świrski, M. I., et al. Translon: a single term for translated regions. Nat. Methods 22, 2002– 2006 (2025).

2. Wright, B. W., Yi, Z., Weissman, J. S. & Chen, J. The dark proteome: translation from noncanonical open reading frames. Trends Cell Biol. 32, 243–258 (2022).

3. Ingolia, N. T., Hussmann, J. A. & Weissman, J. S. Ribosome Profiling: Global Views of Translation. Cold Spring Harb. Perspect. Biol. 11, (2019).

4. Calviello, L. et al. Detecting actively translated open reading frames in ribosome profiling data. Nat. Methods 13, 165–170 (2015).

5. Slavoff, S. A. et al. Peptidomic discovery of short open reading frame–encoded peptides in human cells. Nat. Chem. Biol. 9, 59–64 (2012).

6. Wacholder, A. et al. Detection of human unannotated microproteins by mass spectrometry-based proteomics: a community assessment. bioRxiv. Preprint at 10.1101/2025.02.19.639069. (2025).

7. Vanderperre, B. et al. Direct Detection of Alternative Open Reading Frames Translation Products in Human Significantly Expands the Proteome. PLoS One 8, (2013).

8. Olexiouk, V. et al. sORFs.org: a repository of small ORFs identified by ribosome profiling. Nucleic Acids Res. 44, 324–329 (2016).

9. Mudge, J. M. et al. Standardized annotation of translated open reading frames. Nat. Biotechnol. 40, 994–999 (2022).

10. Sandmann, C. L. et al. Evolutionary origins and interactomes of human, young microproteins and small peptides translated from short open reading frames. Mol. Cell 83, 994–1011 (2023).

11. van Heesch, S. et al. The Translational Landscape of the Human Heart. Cell 178, 242–260 (2019).

12. Chothani, S. P. et al. A high-resolution map of human RNA translation. Mol. Cell 82, 2885– 2899 (2022).

13. Chothani, S. et al. An expanded reference catalog of translated open reading frames for biomedical research. Nucleic Acids Res. 54, (2026).

14. Fesenko, I. et al. A vast pool of lineage-specific microproteins encoded by long non-coding RNAs in plants. Nucleic Acids Res. 49, 10328–10346 (2021).

15. Fesenko, I., Shabalina, S. A., Storz, G. & Koonin, E. V. De novo origin of numerous microproteins in enterobacteria. Nucleic Acids Res. 53, 13–19 (2025).

16. Zhang, L. et al. Rapid evolution of protein diversity by de novo origination in Oryza. *Nat*. Ecol. Evol. 3, 679–690 (2019).

17. Vakirlis, N., Vance, Z., Duggan, K. M. & McLysaght, A. De novo birth of functional microproteins in the human lineage. Cell Rep. 41, (2022).

18. Vakirlis, N. et al. A Molecular Portrait of De Novo Genes in Yeasts. Mol. Biol. Evol. 35, 631–645 (2018).

19. Hassel, K. R., Brito-Estrada, O. & Makarewich, C. A. Microproteins: Overlooked regulators of physiology and disease. iScience 26, (2023).

20. Mohsen, J. J., Martel, A. A. & Slavoff, S. A. Microproteins—Discovery, Structure, and Function. Proteomics 23, (2023).

21. Bornberg-Bauer, E. & Eicholt, L. A. Emergence and evolution of protein-coding de novo genes. Nat. Rev. Genet. 1–17 (2026).

22. Schmitz, J. F., Ullrich, K. K. & Bornberg-Bauer, E. Incipient de novo genes can evolve from frozen accidents that escaped rapid transcript turnover. *Nat*. Ecol. Evol. 2, 1626–1632 (2018).

23. Suenaga, Y. et al. NCYM, a Cis-Antisense Gene of MYCN, Encodes a De Novo Evolved Protein That Inhibits GSK3β Resulting in the Stabilization of MYCN in Human Neuroblastomas. PLoS Genet. 10, (2014).

24. Samusik, N., Krukovskaya, L., Meln, I., Shilov, E. & Kozlov, A. P. PBOV1 Is a Human De Novo Gene with Tumor-Specific Expression That Is Associated with a Positive Clinical Outcome of Cancer. PLoS One 8, (2013).

25. Heames, B. et al. Experimental characterization of de novo proteins and their unevolved random-sequence counterparts. *Nat*. Ecol. Evol. 7, 570–580 (2023).

26. Roginski, P. et al. Impact of GC content on de novo gene birth. Nat. Commun. 17, (2026).

27. Guo, Y. et al. Intrinsic bias of the genetic code shapes the folding and stability landscapes of microproteins. Mol. Cell 86, 2173–2190 (2026).

28. Lange, A. et al. Structural and functional characterization of a putative de novo gene in Drosophila. Nat. Commun. 12, 1667 (2021).

29. Bungard, D. et al. Foldability of a Natural De Novo Evolved Protein. Structure 25, 1687– 1696 (2017).

30. Yamamoto, S. et al. Structural characterization of human de novo protein NCYM and its complex with a newly identified DNA aptamer using atomic force microscopy and small-angle X-ray scattering. Front. Oncol. 13, (2023).

31. Her, C., Yeh, Y. & Krishnan, V. V. The Ensemble of Conformations of Antifreeze Glycoproteins (AFGP8): A Study Using Nuclear Magnetic Resonance Spectroscopy. Biomolecules 9, (2019).

32. Aubel, M. et al. High-throughput Selection of Human *de novo*-emerged sORFs with High Folding Potential. Genome Biol. Evol. 16, (2024).

33. Kumar, M. et al. ELM—the Eukaryotic Linear Motif resource—2024 update. Nucleic Acids Res. 52, 442–455 (2024).

34. Tompa, P., Davey, N. E., Gibson, T. J. & Babu, M. M. A Million Peptide Motifs for the Molecular Biologist. Mol. Cell 55, 161–169 (2014).

35. Davey, N. E., Cyert, M. S. & Moses, A. M. Short linear motifs – ex nihilo evolution of protein regulation. Cell Commun. Signal. 13, (2015).

36. Davey, N. E. et al. Attributes of short linear motifs. Mol. Biosyst. 8, 268–281 (2011).

37. Madhu, P.et al. An Atlas of Short Linear Motif-Mediated Human Protein-Protein Interactions. bioRxiv. Preprint at https://www.biorxiv.org/content/10.64898/2026.07.03.735260v1.full. (2026).

38. Bradburn, D. A. et al. A composite motif in calcimembrin/C16orf74 dictates multimeric dephosphorylation by calcineurin. Nat. Commun. 16, (2025).

39. Xie, L. et al. Biochemistry and Protein Interactions of the CYREN Microprotein. Biochemistry 62, 3050–3060 (2023).

40. Olexiouk, V. & Menschaert, G. Using the sORFs.Org Database. Curr. Protoc. Bioinformatics 65, (2019).

41. Jumper, J. et al. Highly accurate protein structure prediction with AlphaFold. Nature 596, 583–589 (2021).

42. Lin, Z. et al. Evolutionary-scale prediction of atomic-level protein structure with a language model. Science 379, (2023).

43. Muñoz, V. & Serrano, L. Elucidating the folding problem of helical peptides using empirical parameters. Nat. Struct. Biol. 1, 399–409 (1994).

44. Eicholt, L. A., Aubel, M., Berk, K., Bornberg-Bauer, E. & Lange, A. Heterologous expression of naturally evolved putative de novo proteins with chaperones. Protein Science 31, (2022).

45. Lenton, S., Tully, M. D. & Skepö, M. From dilute to concentrated solutions of intrinsically disordered proteins: Sample preparation and data collection. Methods Enzymol. 677, 457– 478 (2022).

46. Jacques, D. A., Guss, J. M., Svergun, D. I. & Trewhella, J. Publication guidelines for structural modelling of small-angle scattering data from biomolecules in solution. Biological Crystallography 68, 620–626 (2012).

47. Jasanoff, A. & Fersht, A. R. Quantitative Determination of Helical Propensities from Trifluoroethanol Titration Curves. Biochemistry 33, 2129–2135 (1994).

48. Buck, M., Radford, S. E. & Dobson, C. M. A Partially Folded State of Hen Egg White Lysozyme in Trifluoroethanol: Structural Characterization and Implications for Protein Folding. Biochemistry 32, 669–678 (1993).

49. Hoshino, M., Hagihara, Y., Hamada, D., Kataoka, M. & Goto, Y. Trifluoroethanol-induced conformational transition of hen egg-white lysozyme studied by small-angle X-ray scattering. FEBS Lett. 416, 72–76 (1997).

50. Goodman, M., Listowsky, I., Masuda, Y. & Boardman, F. Conformational aspects of polypeptides. VIII. Helical assignments via far ultraviolet absorption spectra and optical activity. Biopolymers 1, 33–42 (1963).

51. Culik, R. M., Abaskharon, R. M., Pazos, I. M. & Gai, F. Experimental validation of the role of trifluoroethanol as a nanocrowder. Journal of Physical Chemistry B 118, 11455–11461 (2014).

52. Uversky, V. N. Intrinsically disordered proteins and their ‘Mysterious’ (meta)physics. Front. Phys. 7, (2019).

53. Fehrst, A. Protein Stability. in *Structure and Mechanism in Protein Science: A Guide to Enzyme Catalysis and Protein Folding* (W. H. Freeman, 1999).

54. Myers, J. K., Nick Pace, C. & Martin Scholtz, J. Denaturant m values and heat capacity changes: Relation to changes in accessible surface areas of protein unfolding. Protein Science 4, 2138–2148 (1995).

55. Benz, C. et al. Proteome-scale mapping of binding sites in the unstructured regions of the human proteome. Mol. Syst. Biol. 18, (2022).

56. Davey, N. E., Edwards, R. J. & Shields, D. C. Estimation and efficient computation of the true probability of recurrence of short linear protein sequence motifs in unrelated proteins. BMC Bioinformatics 11, (2010).

57. Nugnes, M. V. et al. DisProt in 2026: enhancing intrinsically disordered proteins accessibility, deposition, and annotation. Nucleic Acids Res. 54, 383–392 (2026).

58. Abramson, J. et al. Accurate structure prediction of biomolecular interactions with AlphaFold 3. Nature 630, 493–500 (2024).

59. Kruse, T. et al. Large scale discovery of coronavirus-host factor protein interaction motifs reveals SARS-CoV-2 specific mechanisms and vulnerabilities. Nat. Commun. 12, (2021).

60. Kussie, P. H. et al. Structure of the MDM2 oncoprotein bound to the p53 tumor suppressor transactivation domain. Science 274, 948–953 (1996).

61. Zondlo, S. C., Lee, A. E. & Zondlo, N. J. Determinants of Specificity of MDM2 for the Activation Domains of p53 and p65: Proline27 Disrupts the MDM2-Binding Motif of p53. Biochemistry 45, 11945–11957 (2006).

62. Bornberg-Bauer, E., Hlouchova, K. & Lange, A. Structure and function of naturally evolved de novo proteins. Curr. Opin. Struct. Biol. 68, 175–183 (2021).

63. Lange, A. et al. Structural and functional characterization of a putative de novo gene in Drosophila. Nat. Commun. 12, (2021).

64. Matsuo, T. et al. Secondary Structure of Human De Novo Evolved Gene Product NCYM Analyzed by Vacuum-Ultraviolet Circular Dichroism. Front. Oncol. 11, (2021).

65. Holehouse, A. S. & Kragelund, B. B. The molecular basis for cellular function of intrinsically disordered protein regions. Nat. Rev. Mol. Cell Biol. 25, 187–211 (2024).

66. Erdős, G., Pajkos, M. & Dosztányi, Z. IUPred3: prediction of protein disorder enhanced with unambiguous experimental annotation and visualization of evolutionary conservation. Nucleic Acids Res. 49, 297–303 (2021).

67. Mirdita, M. et al. ColabFold: making protein folding accessible to all. Nat. Methods 19, 679–682 (2022).

68. Randles, B. M., Pasquetto, I. V., Golshan, M. S. & Borgman, C. L. Using the Jupyter Notebook as a Tool for Open Science: An Empirical Study. arXiv. Preprint at (2018) doi:10.1109/JCDL.2017.7991618. (2018).

69. Meng, E. C. et al. UCSF ChimeraX: Tools for structure building and analysis. Protein Science 32, (2023).

70. Casper, J. et al. The UCSC Genome Browser database: 2026 update. Nucleic Acids Res. 54, 1331–1335 (2026).

71. Blanchette, M. et al. Aligning Multiple Genomic Sequences With the Threaded Blockset Aligner. Genome Res. 14, 708–715 (2004).

72. Harris, R. S. Improved pairwise alignment of genomic DNA. PhD Thesis, Pennsylvania State University (2007).

73. Cowieson, N. P. et al. Beamline B21: high-throughput small-angle X-ray scattering at Diamond Light Source. Biological Crystallography 27, 1438–1446 (2020).

74. Hopkins, J. B. BioXTAS RAW 2: new developments for a free open-source program for small-angle scattering data reduction and analysis. J. Appl. Crystallogr. 57, 194–208 (2024).

75. Schneidman-Duhovny, D., Hammel, M., Tainer, J. A. & Sali, A. FoXS, FoXSDock and MultiFoXS: Single-state and multi-state structural modeling of proteins and their complexes based on SAXS profiles. Nucleic Acids Res. 44, 424–429 (2016).

76. Hajizadeh, N. R., Franke, D., Jeffries, C. M. & Svergun, D. I. Consensus Bayesian assessment of protein molecular mass from solution X-ray scattering data. Sci. Rep. 8, (2018).

77. Piiadov, V., Ares de Araújo, E., Oliveira Neto, M., Craievich, A. F. & Polikarpov, I. SAXSMoW 2.0: Online calculator of the molecular weight of proteins in dilute solution from experimental SAXS data measured on a relative scale. Protein Science 28, 454–463 (2019).

78. Nikolovska-Coleska, Z. et al. Development and optimization of a binding assay for the XIAP BIR3 domain using fluorescence polarization. Anal. Biochem. 332, 261–273 (2004).

