## Supplemental Information for "Human *De Novo* Microproteins are Intrinsically Disordered and Contain Short Linear Motifs"

[1] Department of Medical Biochemistry and Microbiology, Uppsala University, Box 582, Husargatan 3, 751 23, Uppsala, Sweden
[2] Department of Chemistry for Life Sciences, Uppsala University, Box 576, Husargatan 3, 751 23, Uppsala, Sweden
[3] Institute for Evolution and Biodiversity, University of Muenster, Huefferstrasse 1, 48149 Muenster, NRW, Germany
[4] Present address: Protein Engineering and Evolution Unit, Okinawa Institute of Science and Technology Graduate University, Onna Son, Okinawa 904-0495, Japan
[5] Present address: Department of Biochemistry, University of Oxford, Oxford OX1 3QU, United Kingdom
[6] Department of Cell and Molecular Biology, Uppsala University, Box 596, Husargatan 3, 751 24, Uppsala, Sweden
[7] Department of Cell Biology, Charles University, Prague, 12843, Czech Republic
[8] Institute of Organic Chemistry and Biochemistry, Czech Academy of Sciences, Prague, 16000, Czech Republic
[9] Department of Protein Evolution, Max-Planck Institute for Biology, Max-Planck-Ring 5, 72076 Tuebingen, BW, Germany
[10] Science for Life Laboratory, Uppsala University, Uppsala, Sweden.
[#] Corresponding author: Ylva Ivarsson

**Overview of Content:**

**Supplementary Methods**

Computational analysis of motif enrichment

**Supplementary Figures**

**Figure S1.** AlphaFold2 predicted structures of all 30 *de novo* microproteins investigated.

**Figure S2.** ESMFold predicted structures of all 30 *de novo* microproteins investigated.

**Figure S3.** Comparison of predicted structures and sequence properties of the six experimentally investigated *de novo* microproteins.

**Figure S4**. SAXS characterization of the six experimentally investigated *de novo* microproteins.

**Figure S5.** *De novo* microproteins show varying degrees of intrinsic helicity.

**Figure S6.** Motif occurrence and enrichment in *de novo* microproteins and their randomized sequence counterparts.

**Figure S7**. *De novo* microproteins and randomized sequence motifs bind peptide-binding domains.

**Figure S8.** Microprotein peptides bind peptide-binding domains with a broad range of affinities.

**Supplementary Methods Text**

**Supplementary Methods**

**Computational analysis of motif enrichment**

***Motifs and domain clans***

We analysed the consensus regular expressions (RegEx) of the 217 bait domains as manually curated based on consensus motifs defined in the ASHI dataset[1]. Baits with identical RegExs were analysed once, which gave 106 unique RegExs of 2–10 positions. A RegEx consists of literal amino acids, "." for any residue, and bracket classes (e.g. [FW]) that match one residue from the listed set; none of the expressions contains a gap of variable length or an anchor. Every motif therefore has a fixed length $L$ and is described by the sets $S_1, \ldots, S_L \subseteq \mathcal{A}$ of residues allowed at each position, where $\mathcal{A}$ is the alphabet of the 20 standard amino acids. Each bait carries the domain clan assigned in the collection, which gave 34 clans. In addition, each bait has a motif name; a name can point to more than one RegEx, and we therefore also report results for the 88 motif names. The RegExs with their clans and motif names, and all results, are provided in the Supplementary Dataset 3 (**Table S2.a**).

***Sequence preprocessing***

The four sequence sets are described in the Datasets section below. Sequences were split at residues other than the 20 standard amino acids (B, O, U, X, Z), and each uninterrupted stretch of standard residues was treated as a separate sequence; this affected 1 DisProt and 2,229 UniProt sequences. Position bins (below) and observed numbers of sequences with a motif refer to the original sequences. We write $c_n$ for the number of sequences of length $n$ in a set.

***Background statistics***

For each set we counted the amino acids, $C_1(a)$, and the pairs of adjacent residues (dipeptides), $C_2(ab)$, within sequences. The same counts were made separately for five relative position bins: the residue at position $p$ (counted from 0) of a sequence of length $n$ falls into bin $j(p,n) = \min(\lfloor 5p/n \rfloor, 4)$, and a dipeptide is assigned to the bin of its first residue. Together with the distribution of sequence lengths, these counts are the only statistics the null models require. They are computed once per sequence set and stored together with the encoded sequences, so further motifs can be analysed without re-reading the sequence files.

***Null models***

With a pseudocount of $\alpha = 1$, residue frequencies and first-order transition probabilities are

$$\pi(a) = \frac{C_1(a) + \alpha}{\sum_{x \in \mathcal{A}} C_1(x) + 20\alpha}, \qquad T(a \to b) = \frac{C_2(ab) + \alpha}{\sum_{x \in \mathcal{A}} C_2(ax) + 20\alpha}.$$

Both null models were estimated separately for every sequence set:

1. **First-order Markov model (primary).** The first residue of a sequence is drawn from $\pi$, and every following residue from $T$ given the preceding residue. The model reproduces the amino-acid

composition of the set and the preferences between adjacent residues. The residue at position $t$ of a sequence has the distribution $\mu_t = \pi T^{t-1}$.

2. **Composition (IID) model (sensitivity analysis).** Every residue is drawn independently from $\pi$.

The program also computes, for each single RegEx, position-specific versions of both models, in which $\pi$ and $T$ are estimated from the counts of each position bin and applied to the residues in that bin. These were not used for any result reported here.

Because the Markov model is fitted to the dipeptide counts of the same sequences, it reproduces the observed numbers of two-residue motifs (e.g. WL, KR, PL) almost exactly. For the 5 two-residue motifs in our set, fold enrichments under the Markov model were 0.97–1.00 in every set, so for these motifs only the composition model is informative. All *de novo* and random sequences begin with methionine, which neither model reproduces. Most motifs that allow methionine at their first position are therefore more frequent in the first fifth of *de novo* and random sequences than uniform placement predicts. This applies to the only motif enriched in random sequences under the Markov model ([FLM]…W of Pex14), which occurs 1.6 times as often as uniform placement predicts in the first fifth of random sequences, and 1.5 times in *de novo* sequences. Fixing the first residue to methionine in the model is reported as a sensitivity analysis below.

***Expected number of motif occurrences***

Occurrences were counted at every start position, so overlapping occurrences count separately. Under the Markov model, the probability that a motif occurs at a start position whose first residue is $a$ follows from the backward recursion

$$h_L(a) = \mathbb{I}[a \in S_L], \qquad h_i(a) = \mathbb{I}[a \in S_i] \sum_{b \in \mathcal{A}} T(a \to b)\, h_{i+1}(b) \quad (i = L-1, \dots, 1),$$

where $\mathbb{I}[\cdot]$ is 1 if the condition holds and 0 otherwise, and the expected number of occurrences in a set is

$$E = \sum_{n \geq L} c_n \sum_{t=1}^{n-L+1} \sum_{a \in \mathcal{A}} \mu_t(a)\, h_1(a).$$

This is the exact expectation under the model. It equals the result of the automaton-based dynamic programming of our earlier implementation, but requires only one recursion per motif and one cumulative sum over positions per sequence set. Under the composition model, the expression reduces to

$$E = \sum_{n \geq L} c_n\, (n - L + 1) \prod_{i=1}^{L} \pi(S_i), \qquad \pi(S) = \sum_{a \in S} \pi(a).$$

***Expected number of sequences containing a motif***

The probability $P_n(\geq 1)$ that a sequence of length $n$ contains at least one occurrence was computed by forward dynamic programming over a deterministic finite automaton that recognises the motif at any position, with an absorbing state for "motif found". Automaton states were tracked jointly with the last residue, as required by the Markov model. The expected number of sequences with at least one occurrence is $\sum_n c_n P_n(\geq 1)$. Because $\mathbb{E}[X] \geq P(X \geq 1)$ holds for any count $X$, the expected number of occurrences must be at least this value; we verified this for every motif and sequence set under both models as a consistency check between the two calculations.

***Clan enrichment***

For a clan with RegExs $R_1, \dots, R_k$ of lengths $L_1, \dots, L_k$ ($L_{\max} = \max_j L_j$), a sequence position counts as one site if at least one $R_j$ starts there. Overlapping or nested motifs (e.g. [LP]PxY and [ALPV]PxY) are therefore not counted twice. The expected number of sites is

$$E = \sum_n c_n \sum_{t=1}^{n} \sum_{a \in \mathcal{A}} \mu_t\,(a)\, u^{(\min(n-t+1,\, L_{\max}))}(a),$$

where $u^{(m)}(a)$ is the probability that at least one $R_j$ with $L_j \leq m$ matches at a start position whose first residue is $a$. The minimum accounts for windows that extend beyond the end of the sequence. $u^{(m)}$ was computed by forward dynamic programming over the set of RegExs that are still compatible with the residues read so far, stopping at the first complete match. For a single RegEx the expression reduces to the one above. The same calculation was used for the motif names. We verified the calculation by complete enumeration of all sequences of up to five residues and by simulation.

***Observed occurrences***

Each sequence set was scanned for every RegEx. We recorded the number of occurrences (overlapping occurrences counted), the number of sequences with at least one occurrence, and the position bin of the start of every occurrence. Observed clan sites were counted as the union of the start positions of the clan's RegExs.

***Enrichment and statistical significance***

Fold enrichment is the ratio of observed to expected occurrences, $\mathrm{FE} = O/E$. Enrichment was tested with the one-sided Poisson upper tail

$$p = P(X \geq O \mid X \sim \mathrm{Poisson}(E)),$$

i.e. a test for enrichment only. $p$ values were corrected with the Benjamini–Hochberg procedure within each sequence set and null model, across all 106 RegExs (motif level), all 34 clans (clan level) or all 88 motif names; $q \leq 0.05$ was considered significant. Occurrences of motifs that overlap themselves, for example within poly-proline stretches, are not independent, so the Poisson test is anti-conservative for such motifs. We therefore also report how many sequences contain each motif.

***Direct comparison of de novo and random sequences***

Independently of any null model, we compared the numbers of *de novo* and random sequences that contain a motif (or a site of a clan) with a two-sided Fisher's exact test, corrected with the Benjamini–Hochberg procedure across the 106 RegExs or across the 34 clans.

***Comparison of enrichment profiles***

Markov fold enrichments of the 106 RegExs were compared between sequence sets (*de novo* versus random, DisProt versus random, UniProt versus random, *de novo* versus DisProt) with two-sided paired

Wilcoxon signed-rank tests (normal approximation with tie correction; pairs with zero difference were discarded).

#### *Sensitivity analyses of the clan results*

We repeated the clan analysis in three ways. (i) With the alternative RegExs listed for each bait in the collection (column ASHI_regex; 152 unique expressions), *de novo* enrichment of FERM_M, WW, Ubiquitin and SH3 was retained and no other of the nine clans of Fig. 4d became enriched in *de novo* sequences. (ii) Without two-residue motifs, which are uninformative under the Markov model, only the PDZ-like and PKinase clans of Fig. 4d changed. PKinase rose to 1.23-fold in *de novo* sequences ($q = 0.007$), but 17 of its 200 sites start at residue 1 (27 of 181 in random sequences); without these sites the fold enrichment would be about 1.13. (iii) With the first residue fixed to methionine in the Markov model of the *de novo* and random sets, the expected counts of the nine clans of Fig. 4d fell and their fold enrichments rose by 0.01–0.04; no significance call in Fig. 4d changed. Fold enrichments fell (by up to 0.22) only for clans with a motif that allows methionine at its first position. Across all 34 clans, 12 clans were still enriched in *de novo* sequences (10 as before, two lost and two gained), and no clan remained enriched in random sequences. The weak DisProt calls of Fig. 4d depend on how the DisProt regions are prepared (see Datasets): with only identical sequences removed (7,045 sequences), all calls equalled those of the merged set; with all 13,347 records as listed, PKinase (1.04-fold, $q = 0.013$) instead of E-set (1.01-fold, $q = 0.08$) was enriched.

#### *Calibration of the clan test*

For the nine clans of Fig. 4d we simulated 10,000 data sets from the fitted Markov model of the *de novo*, random and DisProt sets, with the observed sequence lengths (random seed 1112068). Simulated means equalled the calculated expectations (ratio 0.999–1.001), and the variance-to-mean ratio of the site counts was 0.96–1.28, so clustering of sites inflates the Poisson test only slightly. Empirical $p$ values from the simulations were below 0.05 for exactly the clans that were significant in the Poisson test. Bootstrapping sequences (10,000 replicates; 2,000 for UniProt; expected counts scaled to the total length of each resample) gave 95% percentile intervals for fold enrichment of 1.17–1.31 (FERM_M), 1.22–1.56 (WW), 1.07–1.17 (Ubiquitin) and 1.08–1.20 (SH3) in *de novo* sequences. An independent count with Python RegExs, using the consensus expressions as written in the collection, reproduced the observed site counts and the numbers of sequences with a site of the nine clans in these three sets.

#### *Positional distribution*

For each motif we compared the occurrences in the five position bins with a uniform distribution over the possible start positions. The expected count in bin $j$ is

$$E_j = O \cdot \frac{W_j}{\sum_{k=1}^{5} W_k},$$

where $W_j$ is the number of start positions in bin $j$ at which a motif of length $L$ fits into a sequence. Deviation was described by the ratio $O_j/E_j$ and Pearson's $\chi^2$ statistic,

$$\chi^2 = \sum_{j=1}^{5} \frac{(O_j - E_j)^2}{E_j},$$

with four degrees of freedom. These values were used only to describe the N-terminal enrichment of motifs that can start with methionine (see Null models).

### Datasets

#### *De novo and random sequences*

To represent recently emerged, putatively *de novo* proteins, we used the sequence library described by Aubel et al.[2], derived from human *de novo*-emerged short open reading frames (sORFs): 3,750 *de novo* sequences of 32–59 residues (155,506 residues in total) and 3,750 random sequences from the same study with identical lengths and similar amino-acid composition. All motif statistics use the complete library, not only the microproteins tested experimentally.

#### *DisProt (annotation release 2025_06)*

To analyse experimentally supported intrinsically disordered regions, we used DisProt annotation release 2025_06 (v9.8). DisProt is a curated resource of proteins with experimentally validated intrinsic disorder. The sequence file has 13,347 records (1.09 million residues), one per annotated region and annotation term (7,114 distinct regions), and regions of the same protein can overlap, so the same residues occur many times. For each DisProt entry (3,201 entries), we therefore placed the residues of all its regions at their annotated positions and joined regions that overlapped or adjoined, so that every annotated residue occurs once. Identical stretches were then kept once (32 removed), and one region with an invalid position (start after end) was kept unchanged. This gave 4,622 sequences (328,749 residues), each analysed as a separate sequence.

#### *UniProt (Swiss-Prot; reviewed)*

As a reference set of curated protein sequences, we used UniProtKB/Swiss-Prot (the reviewed subset of UniProtKB) downloaded in December 2025 (UniProt release 2025_04)[3] . Sequences appearing in DisProt[4] were removed (573,403 sequences remained). Sequences shorter than 10 residues and sequences with more than 30% of residues predicted as disordered by IUPred3[5] (long mode, medium smoothing, residue score $> 0.5$) were removed, leaving 510,643 sequences (181.8 million residues).

### Software

The analysis was run with Python 3.12.3, NumPy 1.26.4 and SciPy 1.11.4. Fisher's exact tests and Wilcoxon signed-rank tests were computed with our own Python implementations; in the tests compared with SciPy 1.11.4 (Fisher's tests of eight clans of Fig. 4d, Wilcoxon test of *de novo* versus random), $p$ values agreed within a relative difference of $10^{-11}$. Result tables and figures were produced with Python 3.13.1, NumPy 2.2.2, openpyxl 3.1.5 and Matplotlib 3.10.0.

### Supplementary Figures

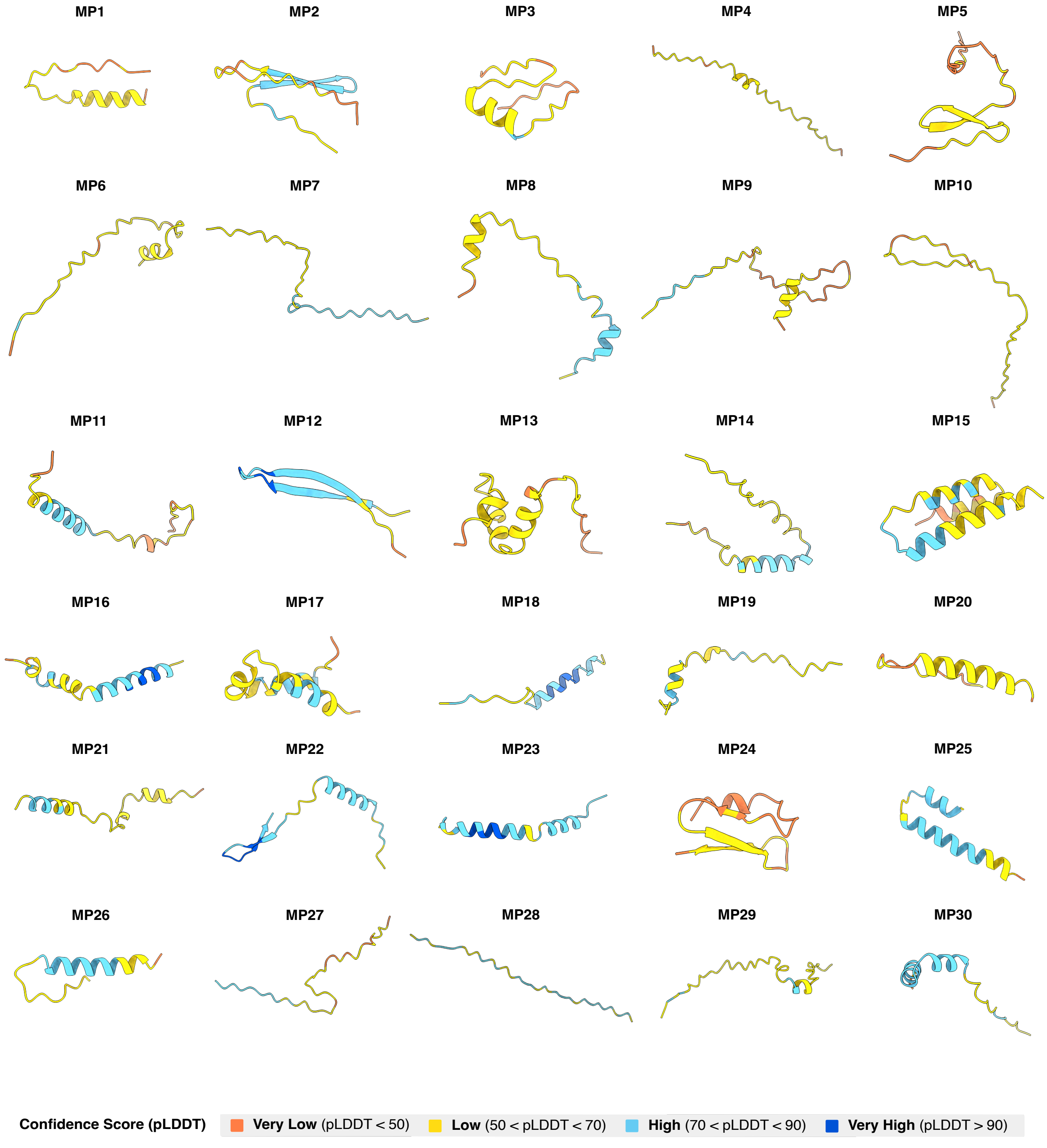


**Figure S1.** AlphaFold2 (AF2)-predicted structures of all 30 *de novo* microproteins investigated. Predicted structures are coloured according to pLDDT scores.

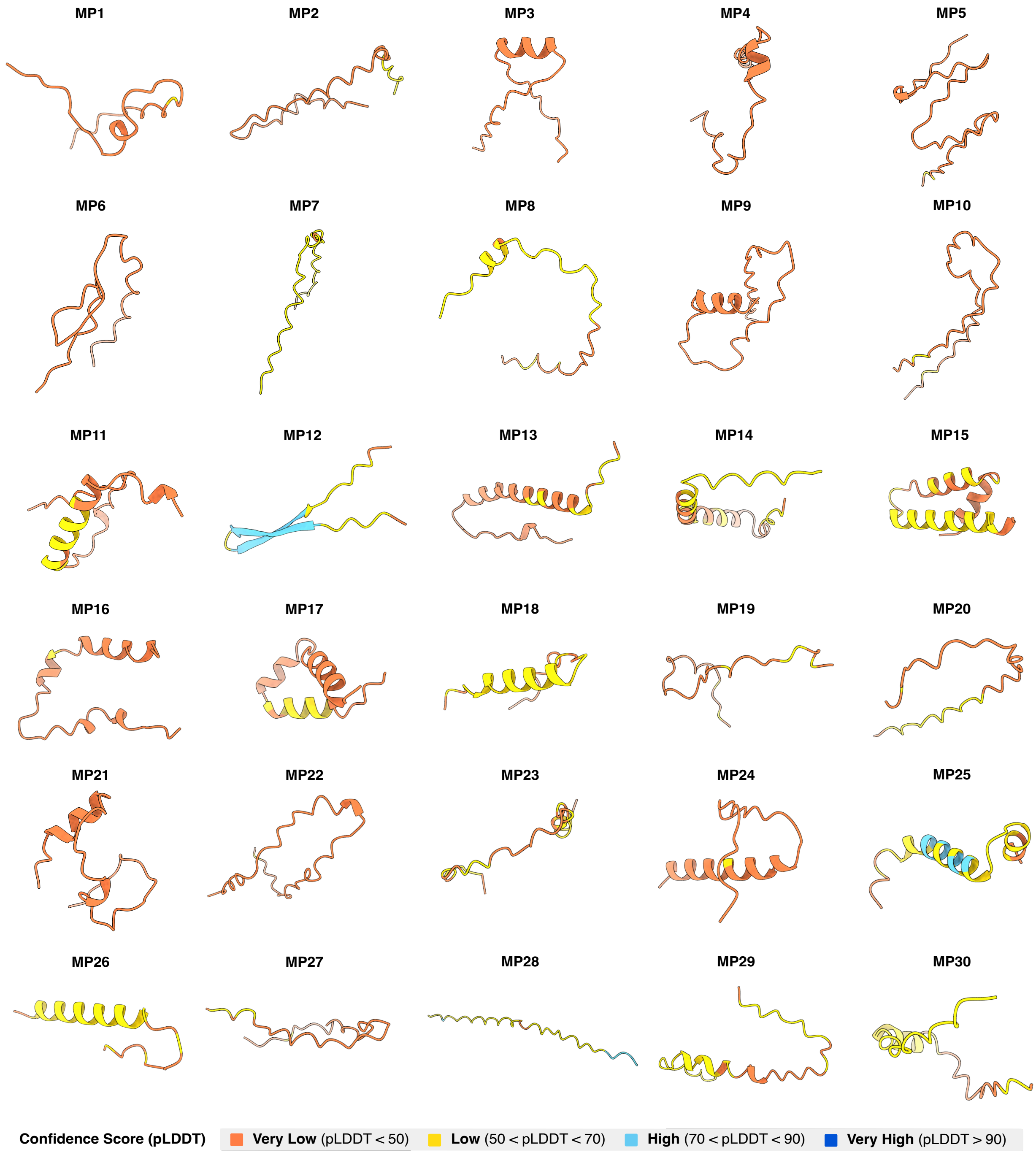


**Figure S2.** ESMFold-predicted structures of all 30 *de novo* microproteins investigated. Predicted structures are coloured according to the pLDDT score.

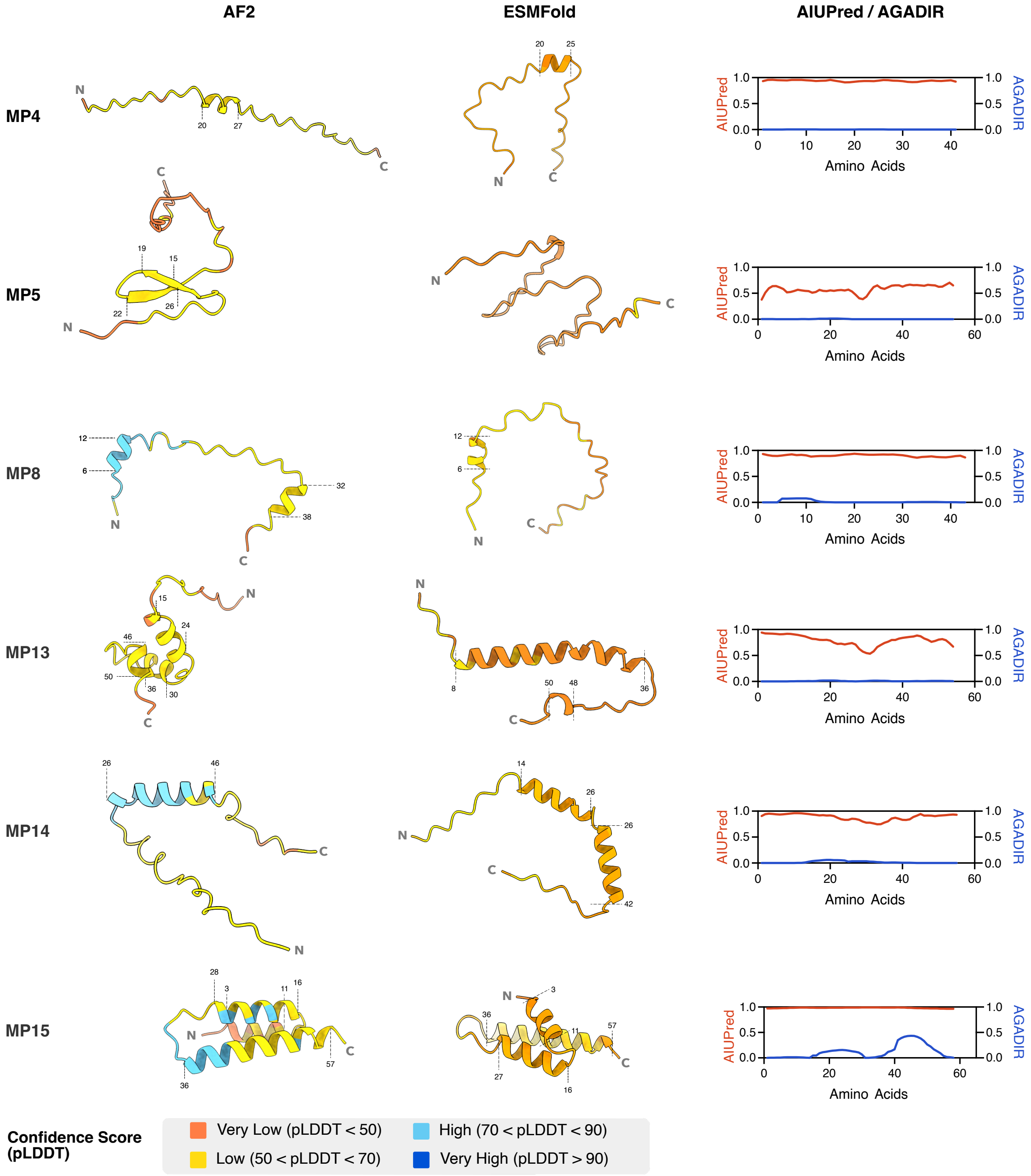


**Figure S3.** Comparison of predicted structures and sequence properties of the six experimentally investigated *de novo* microproteins. Structures predicted with AlphaFold2 (AF2) (left) and ESMFold (middle), with sequence-based predictions of disorder and helicity (right).

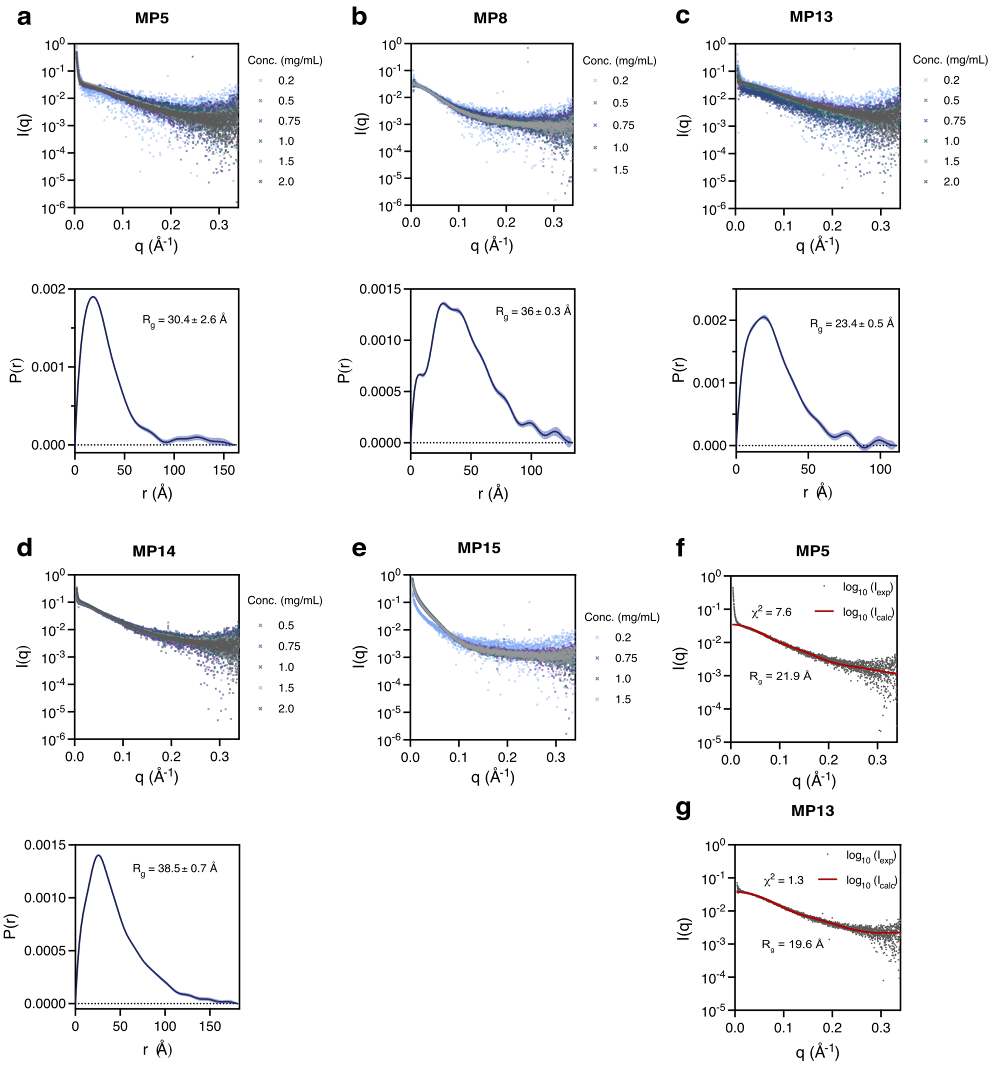


**Figure S4**. SAXS characterization of the six experimentally investigated *de novo* microproteins. Top panels show the experimental scattering curves at all concentrations for (**a**) MP5, (**b**) MP8, (**c**) MP13, (**d**) MP14, and (**e**) MP15. Bottom panels show the corresponding pair distance distribution (P(r)) with the estimated radius of gyration ($R_g$) values. (**f** and **g**) Overlay of experimental (grey) and calculated scattering curves (red) derived from the monomeric structural models of the microproteins. The chi square ($\chi^2$) value indicates the goodness of fit. $R_g$ values estimated from the structural models are given.

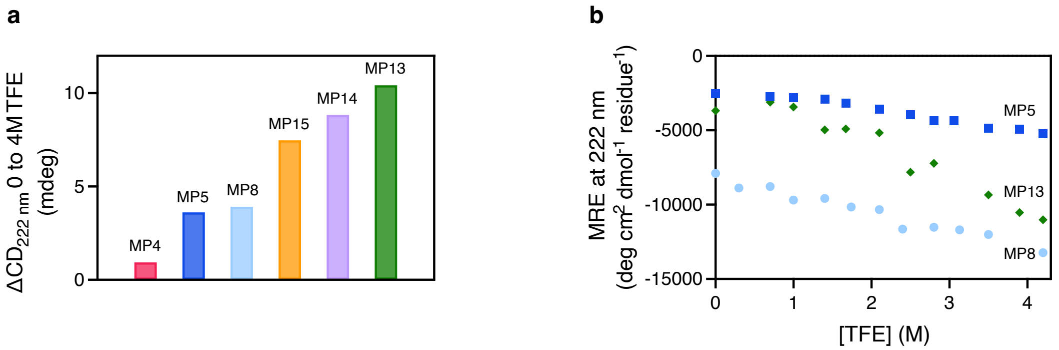


**Figure S5.** *De novo* microproteins show varying degrees of intrinsic helicity. (**a**) Gain in α-helical content, expressed as difference in raw ellipticity (in millidegrees) at 222 nm, upon addition of 4 M TFE in the six experimentally investigated *de novo* microproteins. The microproteins are ranked from lowest (MP4) to highest (MP13) helical propensity. (**b**) Non-sigmoidal transition curves showing the change in mean residue ellipticity (MRE) at 222 nm across the TFE concentration series for MP5, MP8 and MP13. Data are given in MRE, normalized for microprotein size and concentration, to enable comparison between the microproteins.

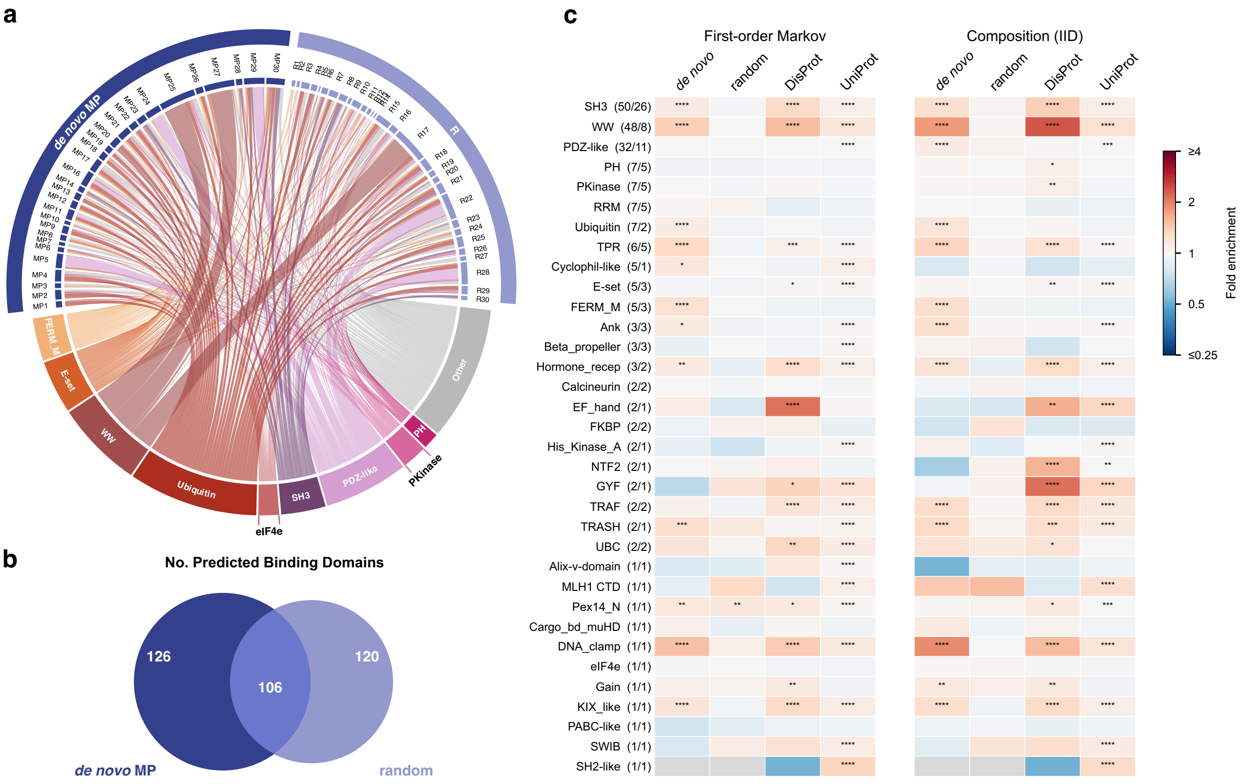


**Figure S6.** Motif occurrence and enrichment in *de novo* microproteins (MP) and their randomized (R) sequence counterparts. (**a**) Chord diagram illustrating all predicted interactions of *de novo* MP sequences and R sequences with all 34 domain clans. Individual chords represent unique MP/R – domain interaction pairs. The thickness of the chords for each clan reflects the total number of unique MP/R – domain interactions, per clan. (**b**) Number of domains binding to predicted motifs in *de novo* MP and/or R sequences. (**c**) Motif enrichment for all 34 bait clans. Fold enrichment of the union of each clan's regular expressions (RegEx) under the first-order Markov (left) and composition (IID, right) null models in the four sequence sets (*de novo*, random, DisProt and UniProt). Numbers in brackets indicate baits per unique RegEx. Stars as in **Figure 4c** (Benjamini-Hochberg over the 34 clans within each set and model). Grey: no site and fewer than 3 expected. Clans that **Table S2** lists as "Other(...)" are shown by their name in brackets

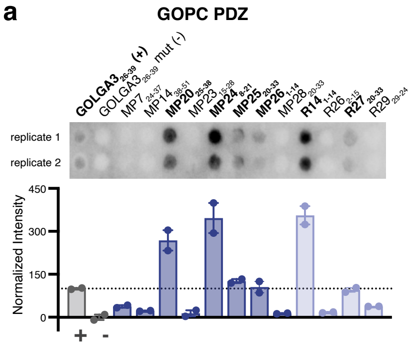


| Protein | Peptide Sequence |
|---|---|
| **GOLGA3$_{26-39}$ (+)** | GLSLPMQETQLCSTDG |
| GOLGA3$_{26-39}$ (-) | GLSLPAQEAQACSTDG |
| MP7$_{24-37}$ | GGPGSLILSPLTAPTG |
| MP14$_{38-51}$ | GLNQQLHPSSLGPKPG |
| **MP20$_{25-38}$** | GNASVDACGLFVSVLG |
| MP23$_{15-28}$ | GLSGPAAGSSLVVAVG |
| **MP24$_{8-21}$** | GYSSRMLNSLLALQIG |
| **MP25$_{20-33}$** | GLLQQLELSILMPYLG |
| **MP26$_{1-14}$** | GMALSSLCSYTVEAFG |
| MP28$_{20-33}$ | GSQFPAPSSHLPAPTG |
| **R14$_{1-14}$** | GMAQTTLTLANRFLGG |
| R26$_{2-15}$ | GIRAAQPTPLCVFHSG |
| **R27$_{20-33}$** | GLSEGWRGTELLSRLG |
| R29$_{29-42}$ | GATRTAAASPLSGGAG |

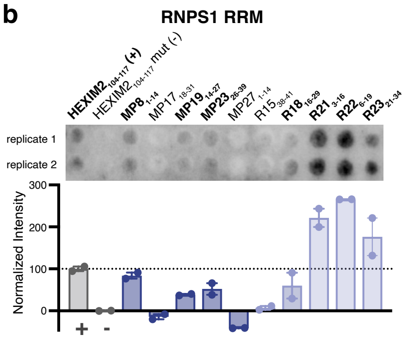


| Protein | Peptide Sequence |
|---|---|
| **HEXIM2$_{104-117}$ (+)** | GPYLELSWAEKQQRDG |
| HEXIM2$_{104-117}$ mut (-) | GPYLEASAASKQQRDG |
| **MP8$_{1-14}$** | GMSWWDPWEAKRSGVG |
| MP17$_{18-31}$ | GPEGKWLAWKHNPLSG |
| **MP19$_{14-27}$** | GMWEPSLFWGVGPTDG |
| **MP23$_{26-39}$** | GVAVVRGLRWLPGLVG |
| MP27$_{1-14}$ | GMKWSDPQGGSDLPQG |
| R15$_{38-41}$ | GGQWFLLLWSPNCLPG |
| **R18$_{16-29}$** | GCAWLPIFWLAELAGG |
| **R21$_{3-16}$** | GRIRELPWRSLFFDGG |
| **R22$_{6-19}$** | GPVDWSELRTVIFWVG |
| **R23$_{21-34}$** | GRNKRLVEWWDSSGSG |

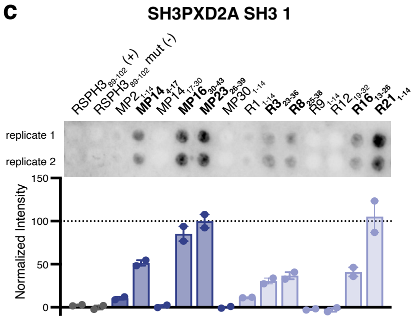


| Protein | Peptide Sequence |
|---|---|
| **RSPH3$_{89-102}$ (+)** | GGNLPSRPPPLLSPLG |
| RSPH3$_{89-102}$ mut (-) | GGNLPSAPAALLSPLG |
| MP2$_{1-14}$ | GMTKYLRLNPTDSALG |
| **MP14$_{4-17}$** | GRPEMPRPRPGREKAG |
| MP14$_{17-30}$ | GAEMEDRRLPYKYAHG |
| **MP16$_{30-43}$** | GAWDLRRRWPFVHANG |
| **MP23$_{26-39}$** | GVAVVRGLRWLPGLVG |
| MP30$_{1-14}$ | GMPPIRTLPLTSPPPG |
| R1$_{1-14}$ | GMHRVYPLQIPPTKGG |
| **R3$_{23-36}$** | GFQHIVRSRPRSRCLG |
| **R8$_{25-38}$** | GWEAYRPAPFLPPSHG |
| R9$_{1-14}$ | GMIARTVPPCASVKLG |
| R12$_{19-32}$ | GLSSGERPVPCITGAG |
| **R16$_{13-26}$** | GPFSKARPEPRRWYGG |
| **R21$_{1-14}$** | GMTRIRELPWRSLFFG |

**Figure S7.** *De novo* microprotein and randomized sequence motifs bind peptide-binding domains. Results of the peptide SPOT arrays for the (**d**) GOPC PDZ, (**e**) RNPS1 RRM, and (**f**) SH3PXD2A SH3 1 domains. Data are represented as in **Figure 4d-f**. Sequences of all peptides (both binding and non-binding) are given on the right.

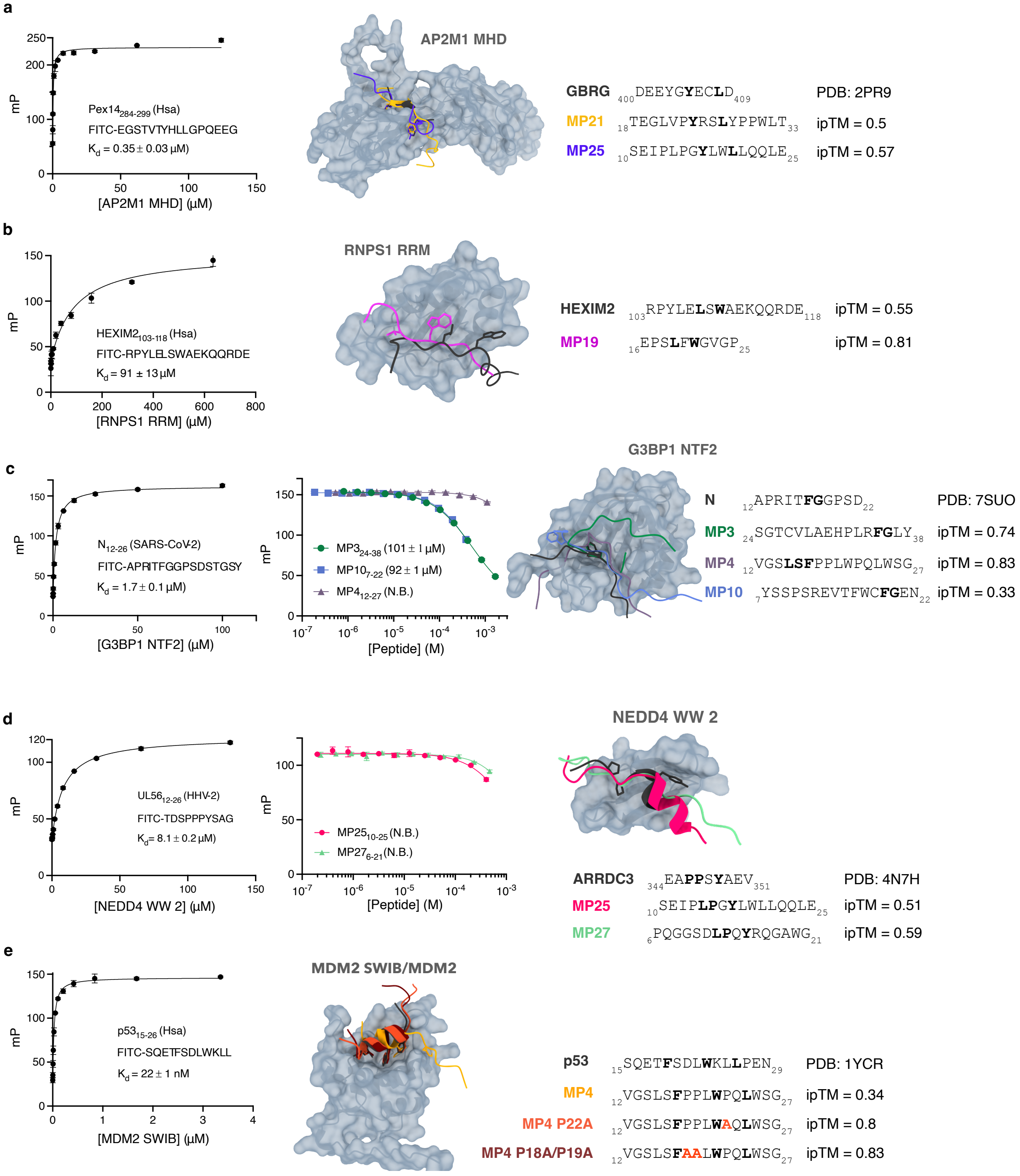
a
Pex14284-299 (Hsa)
FITC-EGSTVTYHLLGPQEEG
Kd = 0.35 ± 0.03 µM
mP
[AP2M1 MHD] (µM)
AP2M1 MHD
GBRG 400DEEYGYECLD409 PDB: 2PR9
MP21 18TEGLVPYRSLYPPWLT33 ipTM = 0.5
MP25 10SEIPLPGYLWLLQQLE25 ipTM = 0.57
b
HEXIM2103-118 (Hsa)
FITC-RPYLELSWAEKQQRDE
Kd = 91 ± 13 µM
mP
[RNPS1 RRM] (µM)
RNPS1 RRM
HEXIM2 103RPYLELSWAEKQQRDE118 ipTM = 0.55
MP19 16EPSLFWGVGP25 ipTM = 0.81
c
N12-26 (SARS-CoV-2)
FITC-APRITFGGPSDSTGSY
Kd = 1.7 ± 0.1 µM
mP
[G3BP1 NTF2] (µM)
MP324-38 (101 ± 1 µM)
MP107-22 (92 ± 1 µM)
MP412-27 (N.B.)
[Peptide] (M)
G3BP1 NTF2
N 12APRITFGGPSD22 PDB: 7SUO
MP3 24SGTCVLAEHPLRFGLY38 ipTM = 0.74
MP4 12VGSLSFPPLWPQLWSG27 ipTM = 0.83
MP10 7YSSPSREVTFWCFGEN22 ipTM = 0.33
d
UL5612-26 (HHV-2)
FITC-TDSPPPYSAG
Kd = 8.1 ± 0.2 µM
mP
[NEDD4 WW 2] (µM)
MP2510-25 (N.B.)
MP276-21 (N.B.)
[Peptide] (M)
NEDD4 WW 2
ARRDC3 344EAPPSYAEV351 PDB: 4N7H
MP25 10SEIPLPGYLWLLQQLE25 ipTM = 0.51
MP27 6PQGGSDLPQYRQGAWG21 ipTM = 0.59
e
p5315-26 (Hsa)
FITC-SQETFSDLWKLL
Kd = 22 ± 1 nM
mP
[MDM2 SWIB] (µM)
MDM2 SWIB/MDM2
p53 15SQETFSDLWKLLPEN29 PDB: 1YCR
MP4 12VGSLSFPPLWPQLWSG27 ipTM = 0.34
MP4 P22A 12VGSLSFPPLWAQLWSG27 ipTM = 0.8
MP4 P18A/P19A 12VGSLSFAALWPQLWSG27 ipTM = 0.83

**Figure S8.** Microprotein peptides bind peptide-binding domains with a broad range of affinities. Affinities of *de novo* microprotein (MP) motifs for their predicted domain binding partners as determined by fluorescence polarization (FP)-monitored competitive binding experiments for (**a**) the AP2M1 MHD domain, (**b**) the RRM domain of RNPS1, (**c**) the NTF2 domain of G3BP1, (**d**) the second WW domain of NEDD4, and the (**e**) SWIB/MDM2 domain of MDM2. Saturation curves (left) were fitted to a quadratic equation to determine the affinities of the domains with their reported ligands. For the G3BP1 NTF2 domain and second WW domain of NEDD4, affinity measurements of non-binding peptides (N.B.) are shown. (right) AlphaFold3 (AF3)-predicted structures of the domains and their respective MP bonders, showing the alignment of the MP peptides and previously reported ligands on the domain binding pockets.

**Supplementary References**


1. Madhu, P. *et al.* An Atlas of Short Linear Motif-Mediated Human Protein-Protein Interactions. *bioRxiv.* Preprint at https://www.biorxiv.org/content/10.64898/2026.07.03.735260v1. (2026).

2. Aubel, M. *et al.* High-throughput Selection of Human *de novo* -emerged sORFs with High Folding Potential. *Genome Biol. Evol.* **16**, (2024).

3. Bateman, A. *et al.* UniProt: the Universal Protein Knowledgebase in 2025. *Nucleic Acids Res.* **53**, 609–617 (2025).

4. Nugnes, M. V. *et al.* DisProt in 2026: enhancing intrinsically disordered proteins accessibility, deposition, and annotation. *Nucleic Acids Res.* **54**, 383–392 (2026).

5. Erdős, G., Pajkos, M. & Dosztányi, Z. IUPred3: prediction of protein disorder enhanced with unambiguous experimental annotation and visualization of evolutionary conservation. *Nucleic Acids Res.* **49**, 297–303 (2021).
